# Efficient Seeding for Error-Prone Sequences with SubseqHash2

**DOI:** 10.1101/2024.05.30.596711

**Authors:** Xiang Li, Ke Chen, Mingfu Shao

## Abstract

Seeding is an essential preparatory step for large-scale sequence comparisons. Substring-based seeding methods such as kmers are ideal for sequences with low error rates but struggle to achieve high sensitivity while maintaining a reasonable precision for error-prone long reads. SubseqHash, a novel subsequence-based seeding method we recently developed, achieves superior accuracy to substring-based methods in seeding sequences with high mutation/error rates, while the only drawback is its computation speed. In this paper, we propose SubseqHash2, an improved algorithm that can compute multiple sets of seeds in one run by defining *k* orders over all length-*k* subsequences and identifying the optimal subsequence under each of the *k* orders in a single dynamic programming framework. The algorithm is further accelerated using SIMD instructions. SubseqHash2 achieves a 10-50*×* speedup over repeating SubseqHash while maintaining the high accuracy of seeds. We demonstrate that SubseqHash2 drastically outperforms popular substring-based methods including kmers, minimizers, syncmers, and Strobemers for three fundamental applications. In read mapping, SubseqHash2 can generate adequate seed-matches for aligning hard reads that minimap2 fails on. In sequence alignment, SubseqHash2 achieves high coverage of correct seeds and low coverage of incorrect seeds. In overlap detection, seeds produced by SubseqHash2 lead to more correct overlapping pairs at the same false-positive rate. With all the algorithmic breakthroughs of SubseqHash2, we clear the path for the wide adoption of subsequence-based seeds in long-read analysis. SubseqHash2 is available at https://github.com/Shao-Group/SubseqHash2.

## 1 Introduction

The ever-growing, gigantic size of sequencing data poses a great challenge for sequence comparison. Seeding is a powerful technique that avoids the expensive computation of all-versus-all, full-length comparisons and is thus widely used in scalable methods. Broadly speaking, seeding transforms a long sequence into a list of shorter ones, often of a regular length, known as seeds. Seeds across sequences can be matched (i.e., compared for identity) in constant time with hash tables. A matching pair of seeds, commonly referred to as a seed-match or anchor, indicates a potential biological relevance and suggests a candidate mapping location. The collected seed-matches may be processed differently to fulfill specific tasks. For example, the seed-chain-extend strategy finds collinear chains of seed-matches to maximize a predefined scoring function. This strategy, often combined with a fine-grained local alignment procedure to fill up the gaps between chained seed-matches, has been widely adopted in read mapping and sequence alignment [2, 3, 18, 1, 9] and is recently shown [26] theoretically to be both accurate and efficient. Another example is identifying all overlapping pairs in a large set of sequences, a critical and the most time-consuming step in constructing the overlap/string graph [10, 19, 6] for genome assembly (following the overlap-layout-consensus paradigm). For this task, seed-matches are used to bucket sequences, thereby confining the search for overlapping pairs within individual buckets and significantly reducing the number of pairs that need to be compared. All these applications demand a seeding scheme that admits both high sensitivity, namely producing many seed-matches on biologically related or similar sequences, and high precision, i.e., producing fewer or ideally zero seed-matches on dissimilar sequences. Both properties are desirable in downstream analyses for producing accurate outcomes; high precision often also implies a reduction of running time.

Substring-based seeding methods, most notably kmers (i.e., substrings of length *k*), have been predominant in sequence analysis, mainly due to their simplicity and extraordinary performance on data with low error rates. However, when comparing sequences with high error rates such as homologous genes from distant species or PacBio and Oxford Nanopore long reads, the choice of *k* is usually full of compromises and frustrations. Using a large *k* guarantees high precision but is prohibitive for its extremely low sensitivity. Conversely, a small *k*, improves sensitivity, but because unrelated sequences may share many short kmers by chance, it often results in poor precision. This inherent dilemma of kmers makes it much less effective when the mutation/error rate is high. Many tools are forced to use a small *k* and compensate for the low precision by fine-grained follow-up steps such as chaining to filter out false positives, which significantly increases processing time and highlights the deficiency of using kmers for seeding. Existing sketching methods such as minimizers [25, 22, 21, 16] and syncmers [8] can reduce the number of seeds and seed-matches but cannot solve the sensitivity-precision perplexity of kmers. (See the comparisons of kmers, minimizers, and syncmers in Figures 4, 5, and 6.) Spaced seeds [5, 13] and indel seeds [15] accommodate errors by masking positions with specific patterns, but only mutations at designated positions can be handled. Another strategy that recently gains popularity is to combine multiple short kmers to produce a longer seed; examples include neighboring minimizer pairs [7], Order Min Hash [17], and Strobemers [23, 14, 24]. Nonetheless, as these methods are all substring-based, they cannot fully resolve the intrinsic weakness of kmers against mutations. Additionally, sketching and combining can be considered orthogonal to the basic seeding methods in the sense that they can be applied with any basic seeding scheme (see Section 2.5).

In [12], we proposed a novel subsequence-based seeding approach named SubseqHash. The key intuition is that two strings have a small edit distance *if and only if* they share long subsequences. More concretely, two length-*n* strings with an edit distance of *e* must share some subsequence of length at least *n* − *e*, and two length-*n* strings sharing a subsequence of length *k* must admit an edit distance at most 2(*n* − *k*). In contrast, two length-*n* strings with an edit distance of *e* can only guarantee to share a substring/kmer of length *n* − *ke*, as a single edit can break *k* continuous kmers. SubseqHash is formally defined as a function *h_π_*(***x***) that maps a length-*n* string ***x*** to the smallest length-*k* subsequence of ***x*** according to *π*, which is an order over all length-*k* strings/subsequences. Note that the number of subsequences of ***x*** grows exponentially; we made an algorithmic breakthrough to overcome this: we designed a specific order *π* termed the ABC order, along with an algorithm to calculate *h_π_*(***x***) under an ABC order *π* in *O*(*nkd*) time where *d* is a parameter that balances the probability of hash collision and the running time. We demonstrated that the probability of hash collision Pr(*h_π_*(***x***) = *h_π_*(***y***)) is close to the Jaccard similarity of two sets of subsequences, defined as *J_k_*(***x***, ***y***) := *|S_k_*(***x***)∩*S_k_*(***y***)*|/|S_k_*(***x***)∪*S_k_*(***y***)*|* where *S_k_*(*·*) denotes the set of length-*k* subsequences in a string. SubseqHash inherently tolerates errors/edits by design, making it particularly suitable for seeding sequences with high error rates.

Although SubseqHash achieved better accuracy than substring-based methods on data with high error rates, there is still room for improvement. Specifically, a single run of SubseqHash cannot achieve both high sensitivity and high precision. To control false positives, we typically choose a large *n* (e.g., *n* = 30, 40, 50) and a *k* that is close to *n* (e.g., *k* = 0.7*n,* 0.8*n,* 0.9*n*), as dissimilar strings are not expected to share subsequences of that long and hence a hash collision will not happen, resulting in the desired high precision. But with such *n* and *k*, the probability of hash collision *p* := Pr(*h_π_*(***x***) = *h_π_*(***y***)) is not high enough for similar strings ***x*** and ***y***, since the portion of shared length-*k* subsequences measured with *J_k_*(***x***, ***y***) is small, leading to low sensitivity. To address this, we proposed a new seeding scheme by *repeating* SubseqHash multiple times (say *t* times), with independent and random ABC orders. Suppose that *p* is the probability of a hash collision when invoking SubseqHash once, the probability of having at least one hash collision among the *t* pairs of seeds can be boosted to 1 − (1 − *p*)*^t^*. Note that for similar strings, *p* is strictly positive (thanks to the fact that similar strings share long subsequences), thus sensitivity is markedly enhanced. Conversely, dissimilar strings, which are unlikely to share any long subsequence, will still have a zero or negligibly small probability of hash collision, implying minimal precision degradation with repeats. SubseqHash combined with repeating *can* yield both high sensitivity and precision, as we experimentally demonstrated in Section 3.1 and in the SubseqHash paper [12].

However, the above seeding scheme is computationally inefficient. In practice, sliding windows of length-*n* are used on long reads and SubseqHash is repeatedly apply to each window. Let *N* be the length of a long read and hence it contains *N* − *n* + 1 = *O*(*N*) length-*n* windows. The overall running time of seeding for this single long read is *O*(*Nnkdt*), which could be prohibitive for certain applications. This computational demand limits the widespread use of SubseqHash. We address this challenge in this paper by introducing SubseqHash2, which seeds for a length-*N* long read with *t* repeats in *O*(*Nnk*) time, a *dt*-fold speedup over SubseqHash. The acceleration is achieved by two algorithmic innovations. First, we design the ABCk orders, which defines *k* orders, and a new dynamic programming algorithm that finds the optimal subsequences for the *k* orders simutaneously. A single run of this algorithm generates *k* seeds, leading to a *t*-fold speedup over SubseqHash for any *t* ≤ *k*. Second, recognizing the independence of the subproblems in the dynamic programming algorithm, we leverage SIMD (single instruction, multiple data) parallelism that solves *d* subproblems in parallel with one set of instructions. This gains another *d*-fold speedup, for *d* up to 32. In practice, SubseqHash2 may not experience exactly a *dt*-fold speedup due to the overheads, but the acceleration scales with *dt*. For example, with *n* = 30 and a typical choice of *k* = 25 and *d* = 31 the observed speedup is about 50*×*.

In addition to the improved efficiency, the design of SubseqHash2 allows for symmetries in the score function, enable a string and its reverse complement to produce the same sets of seeds. This feature is highly desireable in sequence analysis, as it effectively halves both running time and storage requirements. We also show that SubseqHash2 can be extended to handle multiple windows and be combined with kmers. SubseqHash2 is now ready for practical use, as demonstrated in three applications including read mapping, pairwise alignment, and overlap detection. In all experiments, SubseqHash2 obtains much higher accuracy compared with substring-based seeding methods; it consistently mirrors the seed quality and accuracy of SubseqHash but with a substantial reduction in running time.

## 2 SubseqHash2

The key idea of SubseqHash2 is the introduction of a pivot position that bifurcates the length-*k* subsequence into two disjoint subsequences. For each pivot, a score function can be defined by amalgamating the scores of the two subsequences and the pivot position, resulting in *k* score functions (i.e., *k* orders). Intriguingly, the optimal subsequences under the *k* orders can be computed together in a single dynamic programming framework, which takes 1*/k* running time compared to SubseqHash. The algorithm is further accelerated by SIMD, which solves *d* subproblems simutaneously, leading to a *d*-fold speedup. We also introduce the two variants of SubseqHash2: SubseqHash2r and SubseqHash2w.

### 2.1 The ABCk Orders

We define the *k* orders over Σ*^k^*, termed the ABCk orders. (See an example in Appendix B) They are governed by an integer *d* and 9 tables: forward tables *A_F_, B_F_, C_F_*, reverse tables *A_R_, B_R_, C_R_*, and pivot tables *A_P_, B_P_, C_P_*. Tables *A_F_* and *A_R_* are of dimension *k × d × |*Σ*|*, i.e., *A_F_, A_R_* ∈ *Z*^*k×d×|Σ|*^. Table *A_P_* is of dimension *k × |*Σ*|*, i.e., *A_P_* ∈ ℤ^*k*×|*Σ*|^. Tables *B_F_* and *B_R_* are of dimension *k × d × |*Σ*|*, and table *B_P_* is of dimension *k × |*Σ*|*; each element in them is a pair, namely, *B_F_* [*i*][*j*][*σ*]*, B_R_*[*i*][*j*][*σ*]*, B_P_* [*i*][*σ*] ∈ {(+1, +1), (+1, −1), (−1, +1), (−1, −1)}, 1 ≤ *i* ≤ *k*, 0 ≤ *j* ≤ *d* − 1, and *σ* ∈ Σ. Tables *C_F_*, *C_R_* and *C_P_* are of dimension *k × |*Σ*|*, where *C_F_* [*i*][*σ*]*, C_R_*[*i*][*σ*]*, C_P_* [*i*][*σ*] ∈ {0, 1*, …, d* − 1}, 1 ≤ *i* ≤ *k* and *σ* ∈ Σ. These tables can be filled either by picking values from their specific ranges randomly, or in a symmetric way for a desired property (Section 2.4).

These nine tables, once filled, determine *k* orders over Σ*^k^*, named *π*_1_*, π*_2_*, …, π_k_*. All these orders utilize forward functions *ψ_F_* and *ω_F_*, as well as reverse functions *ψ_R_* and *ω_R_*, which we describe now. The three forward tables are used in the forward functions while the three reverse tables are used in the reverse functions. Let ***s*** ∈ Σ*^l^* be a string of length *l*, where *l* ≤ *k*. Write ***s*** = *s*_1_*s*_2_ *… s_l_*. Denote the first and the second element in the pair *B_F_* [*i*][*j*][*σ*] by *B_F_* [*i*][*j*][*σ*]_1_ and *B_F_* [*i*][*j*][*σ*]_2_, respectively; the same applies for the tables *B_R_* and *B_P_*. Then the functions are defined by the following recurrences:

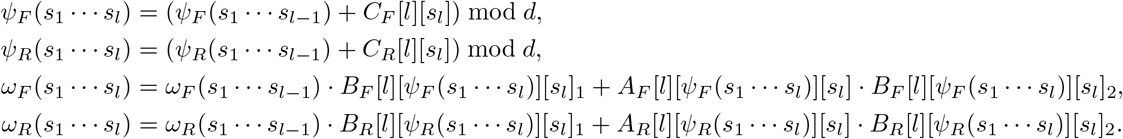

The initial values are set to *ψ*_*F*_ (***s***) = *ψ R*(***s***) = 0 and *ω*_*F*_ (***s***) = *ω*_*R*_ (***s***) = 0 for empty string ***s***.

We now define the *k* orders *π*_*i*_, *i* = 1, 2*, …, k*, over Σ*^k^*. Each order *π*_*i*_ is essentially a score function that maps a length-*k* string to a pair. Let ***z*** = *z*_1_*z*_2_ *… z_k_* ∈ Σ*^k^*. Intuitively, *π*_*i*_ picks *z_i_* as the pivot and combines the forward function and the reverse function using the pivot tables. Formally, we define *π*_*i*_(***z***) := (*ψ*_*i*_(***z***), *ω*_*i*_(***z***)), where

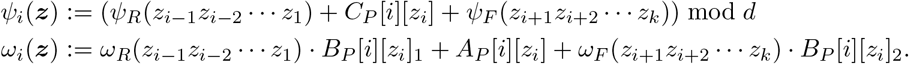

For any two ***z*_1_**, ***z*_2_** ∈ Σ*^k^*, we define *π*_*i*_(***z*_1_**) *< π*_*i*_(***z*_2_**), i .e., ***z* _1_** is ranked before ***z* _2_** in order *π*_*i*_, if and only if*ψ _i_*(***z*_1_**) *<ψ _i_*(***z*_2_**), or *ψ* _*i*_(***z*_1_**) =*ψ _i_*(***z*_2_**) and *ω*_*i*_(***z*_1_**) *> ω*_*i*_(***z*_2_**).

Similar to the ABC order introduced in [12], the techniques of using *±*1 in *B_F_*, *B_R_* and *B_P_* tables, the modular operation, and the pivot splitting all aim for assigning shuffled scores to two strings with small edit distance, so that similar strings are distant in the resulting order. This is critical in achieving a higher probability of hash collision. Note also that the *k* orders in the ABCk orders are not independent as they use shared forward and reverse functions, but we experimentally show that the *k* orders behave independently in practice, and SubseqHash2 achieves almost identical results with repeating SubseqHash using independent tables (see Section 3).

### 2.2 Algorithm for Computing Seeds

A common practice in handling (long) sequences is to generate seeds for each of its sliding window (i.e., length-*n* substring). Here, we design an algorithm that finds seeds for all windows in a given long sequence at one time, which runs Θ(*k*) times faster than processing each sliding window separately. Let ***X*** be a sequence of length *N*, *N > n*. We use *X*[*w|b*] to denote the length-*b* substring of *X* starting from position *w*, i.e., *X*[*w|b*] := *X*_*w*_*X*_*w*+1_ … *X*_*w*+*b*−1_. We define

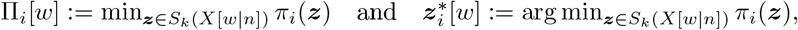

for every 1 ≤ *i* ≤ *k* and 1 ≤ *w* ≤ *N* − *n* + 1. The set 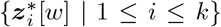 gives the *k* optimal seeds for window *X*[*w|n*], corresponding to the *k* orders {*π_i_ |* 1 ≤ *i* ≤ *k*}, while {*Π*_*i*_[*w*] *|* 1 ≤ *i* ≤ *k*} gives the optimal scores.

Given ***X***, we design an algorithm to calculate Π*_i_*[*w*] and 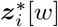. As shown in Figure 1, the algorithm consists of two steps, *iterating* and *optimizing*, following a typical dynamic programming scheme. In the iterating step, subproblems are defined and solved with recurrences. In the optimizing step, the smallest subsequences under each order for every window are calculated. Different windows may reuse the same subproblems, making this algorithm Θ(*k*) times faster than processing windows separately.

**Figure 1.**
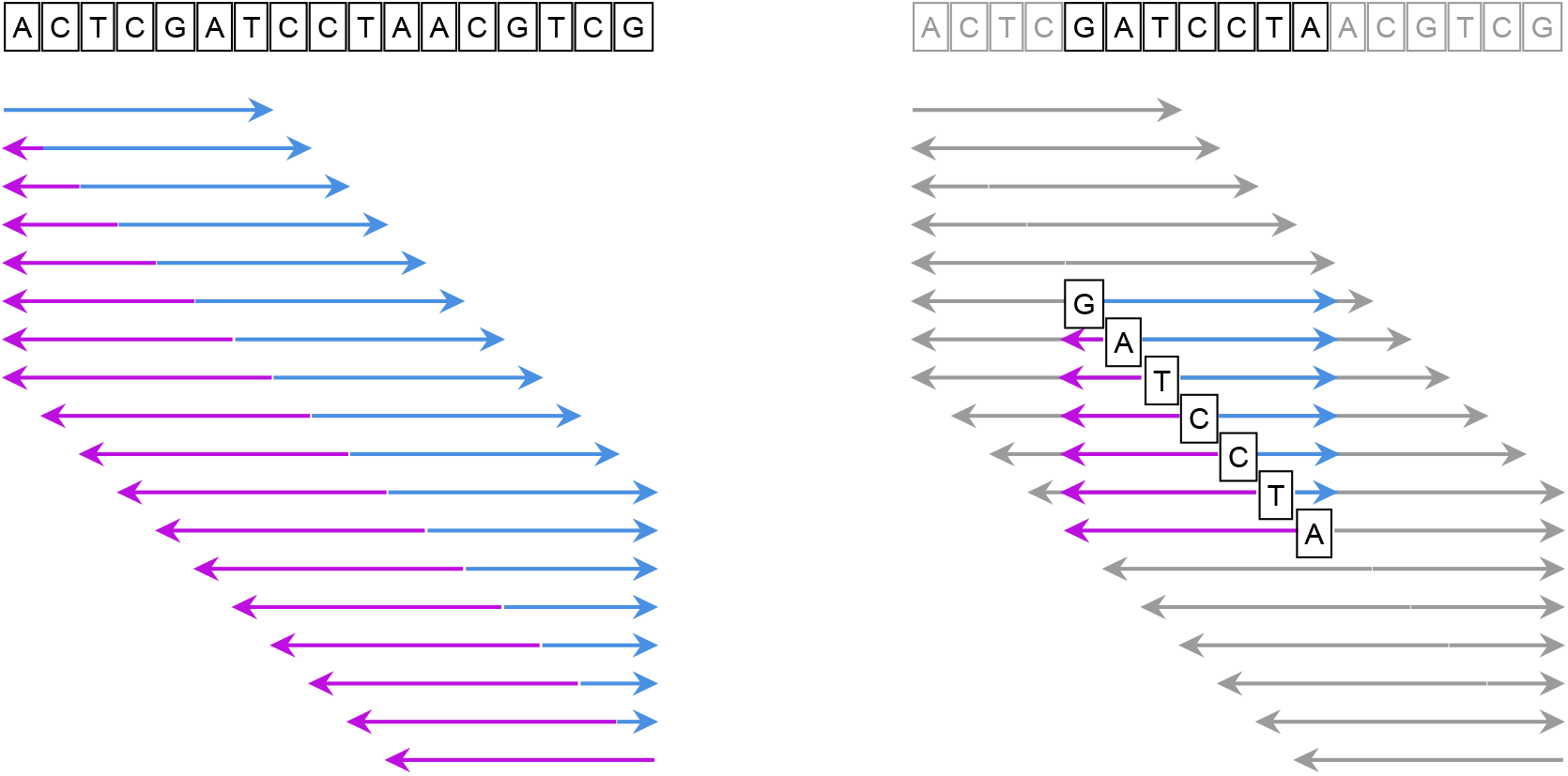
Illustration of the algorithm computing seeds for each window of length *n* = 7 of the given long sequence (on top; length *N* = 17). The left panel shows the iterating step, in which the blue arrows show the *F*_*min*_ and *F*_*max*_ subproblems and purple arrows show the *R*_*min*_ and *R*_*max*_ subproblems. The right panel shows the optimizing step for a particular window (highlighted); the subproblems required for finding the optimal seeds in this window, which are available in the iterating step, are also highlighted.

For any string ***x***, we use *S_l_*(***x***) to denote the set of length-*l* subsequences of *x, l* ≤ *|****x****|*. For each 1 ≤ *w* ≤ *N* − *n* + 1, 1 ≤ *b* ≤ *n*, 1 ≤ *l* ≤ *k*, and 0 ≤ *j < d*, we define subproblem *F* _min_[*w*][*b*][*l*][*j*], where *w* and *b* specify window *X*[*w|b*], *l* indicates the length-*l* subsequences of *X*[*w|b*], *j* restricts that the ψ value of the subsequences must be *j*, and *F*_min_[*w*][*b*][*l*][*j*] is defined as the smallest *ω* value among all such subsequences. Similarly we can also define subproblems *F* _max_[*w*][*b*][*l*][*j*]. Their formal definitions are given below.

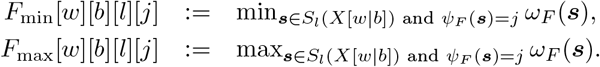

We also define subproblems *R*_min_[*w*][*b*][*l*][*j*] and *R*_max_[*w*][*b*][*l*][*j*], which are to compute the minimized and maximized *ω* value among all *reversed* length-*l* subsequences in window *X*[*w|b*] given their *ψ*value must be equal to *j*. Their definitions are given below, in which we use 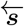 to represent the reversed string of *s* (i.e., if ***s*** = *ACGAT* then 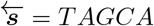).

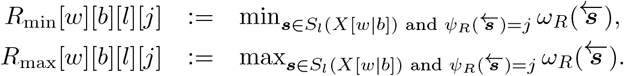

The iterating-steps solves all above subproblems. Since both *ψ* and *ω* functions are recursively defined, it is straightforward to give the recurrences. Specifically, for *F*_min_ and *F*_max_ we consider if the last (the *l*-th) letter of the optimal *s* comes from the the (i.e. *X*_*w*+*b*− 1_) letter of the window. Whether *F_min_* or *F*_max_ gets used for the smaller subproblem depends on the binary vector in the corresponding *B*_*F*_ table. The recurrences are given below, in which we define *σ* := *X*_*w*+*b*−1_ and *j*′ := (*j* − *C_F_* [*l*][*σ*] + *d*) mod *d*:

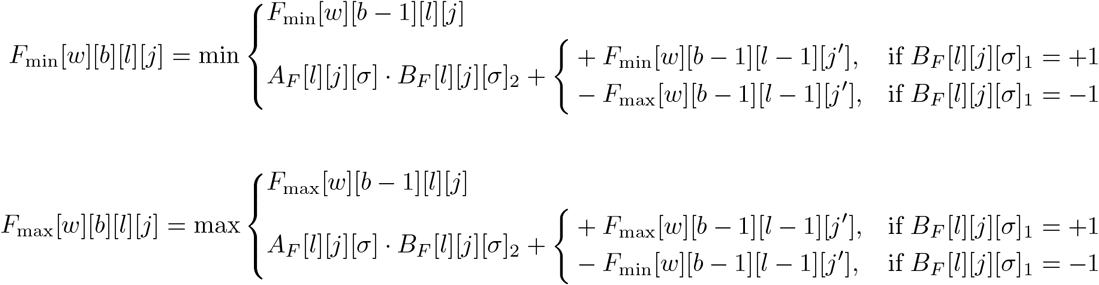

For *R*_min_ and *R*_max_ we consider whether the last letter of the optimal 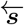, which is the first letter of ***s***, comes from the first letter of the window. Define *σ* := *X_w_* and *j*′ := (*j* − *C_R_*[*l*][*σ*] + *d*) mod *d*. We have

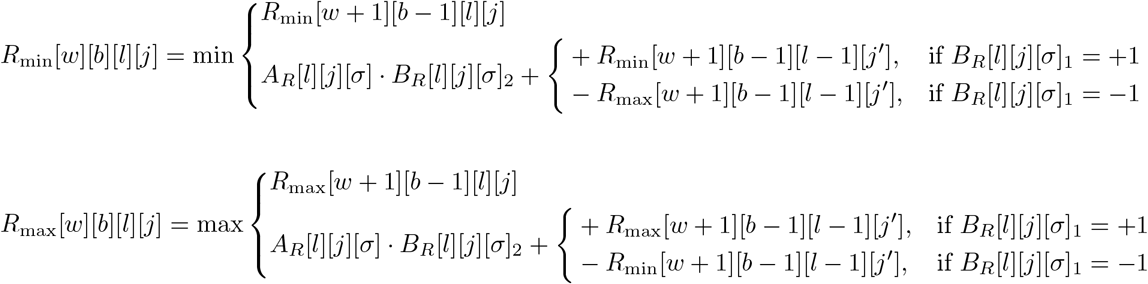

Initially, for any 1 ≤ *w* ≤ *N* − *n* + 1 and 1 ≤ *b* ≤ *n, F_min_*[*w*][*b*][0][0] = *F_max_*[*w*][*b*][0][0] = 0, *R_min_*[*w*][*b*][0][0] = *R_max_*[*w*][*b*][0][0] = 0, *F_min_*[*w*][*b*][0][*j*] = *F_max_*[*w*][*b*][0][*j*] = 0, and *R_min_*[*w*][*b*][0][*j*] = *R_max_*[*w*][*b*][0][*j*] = NaN if *j* ≠ 0. When applying above recurrences to solve all subproblems, if any of the arithmetic operations {+,−} involves NaN as an operand, then the result is also an NaN. The min and max operations ignore NaN and only work on numerical operands, unless there is none, in which case an NaN is returned. We essentially use NaN to indicate that no feasible subsequence exists for a subproblem. There are *O*(*Nnkd*) subproblems defined; solving all of them using above recurrences takes *O*(*Nnkd*) time.

The optimizing-step calculates the optimal score 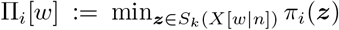 and the optimal subsequence 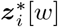 under each order *π*_*i*_ for each window *X*[*w|n*] by using the solutions of above subproblems. We enumerate where the *i*-letter of 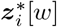, which is the pivot position for order *π*_*i*_, comes from in *X*[*w|n*]. Having, say *X*_*w*+*b*−1_ is the *i*-the letter for a particular *b*, 1 ≤ *b* ≤ *n*, then calculating Π*_i_*[*w*] becomes two smaller problems involving finding a length-(*i* − 1) subsequence ***s*_1_** from *X*[*w|b* − 1] and a length-(*k* − *i*) subsequence ***s*_2_** from *X*[*w* + *b|n* − *b*]. Formally, we write

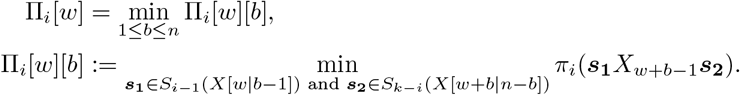

Recall that *π*_*i*_(*·*) := (*ψ_i_*(*·*)*, ω_i_*(*·*)), Therefore we need to find the optimal *ψ_i_* and *ω_i_*, defined as

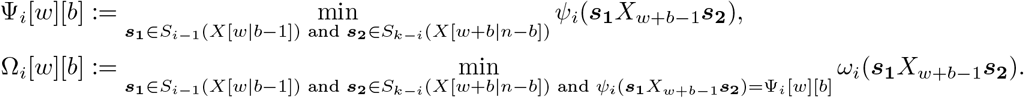

We design an *O*(*d*) time algorithm to calculate Ψ*_i_*[*w*][*b*]. Recall the definition of*ψ_i_* a nd n ote that *X*_*w*+*b*−1_ is the pivot, we have

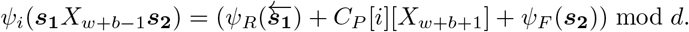

We first collect values 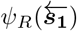 and *ψ_F_* (***s*_2_**) can take; we use two vectors, *V* _1_ and *V*_2_, to store t hem. For each *j* = 0, 1*, …, d* − 1, we define *j* ∈ *V* _1_ if and only if there exists 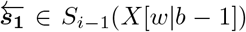 such that 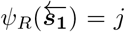; similarly, we define *j* ∈ *V*_2_ if and only if there exists ***s_2_*** ∈_*k*−*i*_ *S* (*X*[*w* + *b|n* − *b* ]) such that *ψ_F_*(***s*_2_**) = *j*. These two (sorted) vectors can be calculated easily given the solutions of the subproblems in the iterating step. In fact, *j* ∈ *V*_1_ if and only if *R*_min_[*w*][*b* − 1][*i* − 1][*j*] ≠ NaN, and *j* ∈ *V*_2_ if and only if *F*_min_[*w* + *b*][*n* − *b*][*k* − *i*][*j*] */*= NaN. Therefore Ψ*_i_*[*w*][*b*] can be calculated as:

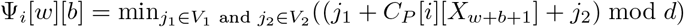

This optimization problem can be solved in *O*(*d*) time, assuming that a binary vector of size *d* can be fit in a machine word, or in *O*(*d ·* log *d*) time for arbitrary *d*. We leave the algorithmic details in Appendix A.

As for *Ω*_*i*_[*w*][*b*], to satisfy that *ψ_i_*(***s*_1_***X*_*w*+*b*−1_***s*_2_**) = Ψ*_i_*[*w*][*b*], we enumerate how it gets distributed across the three parts: if we assume that 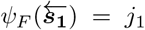, then we know *ψ_R_*(***s_2_***) = (Ψ*_i_* [*w*][*b*] − *j*_1_ −*C_P_* [*i*][*X*] _*w*+*b*−1_ ]+ *d*) mod *d,* which we define as *j*_2_. Recall that

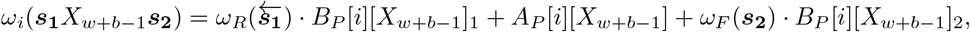

we have

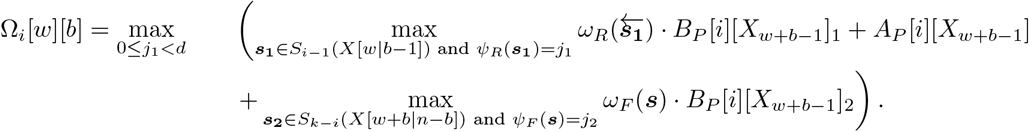

The two optimization problems here are already solved in the iterating-step: the binary values in *B_P_* [*i*][*X*_*w*+*b*−1_] determines whether *F*_min_/*R*_min_ or *F*_max_/*R*_max_ should be used. If we define

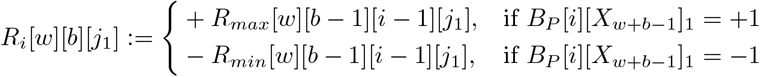

and

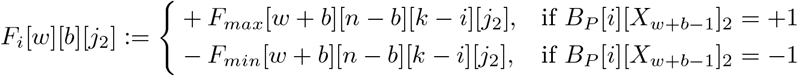

Then *R_i_*[*w*][*b*][*j*_1_] and *F_i_*[*w*][*b*][*j*_2_] exactly give the optimal solution of the first and third term in the parenthesis. Formally,

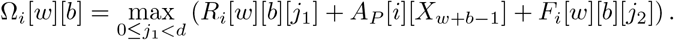

Calculating Ω*_i_*[*w*][*b*] and thus Π*_i_*[*w*][*b*] = (Ψ*_i_*[*w*][*b*], Ω*_i_*[*w*][*b*]) takes *O*(*d*) time for a single pair of *w* and *b*. Hence, the total running time of optimizing-step is *O*(*Nnkd*).

In the end, Π*_i_*[*w*] is computed as min_1≤*b*≤*n*_ Π*_i_*[*w*][*b*], and the optimal subsequence 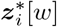 can be obtained by traceback. Therefore, the entire algorithm that finds *k* seeds for every length-*n* window in a given length-*N* sequence also runs in *O*(*Nnkd*) time.

### 2.3 SIMD Parallelism

The total running time of the above algorithm is *O*(*Nnkd*), as both iterating-step and optimizing-step require either filling or tracking back of a dynamic programming table of dimension *N × n × k × d*. Here we use SIMD instructions to speedup the algortihm. SIMD instructions are widely supported by modern CPUs that can leverage wide registers to perform operations on multiple data elements in parallel. Many dynamic programming algorithms are not feasible to use SIMD to speedup as the subproblems usually depend on each other. In our case, the subproblems in the last dimension of the table can be made independent, and hence these subproblems can be solved together with SIMD instructions.

The iterating-step solves *O*(*Nnkd*) subproblems in *O*(*Nnkd*) time. With SIMD parallelism, we can concurrently compute *d* subproblems. Within the recurrences, operations such as addition, multiplication, minimum, and maximum operations can be replaced with SIMD instructions in the CPU intrinsics instruction set. Take the subproblem *F*_min_[*w*][*b*][*l*][*j*] as an example, we can get all the values of *F*_min_[*w*][*b*][*l*][*j*], 0 ≤ *j < d* in *O*(1) time.

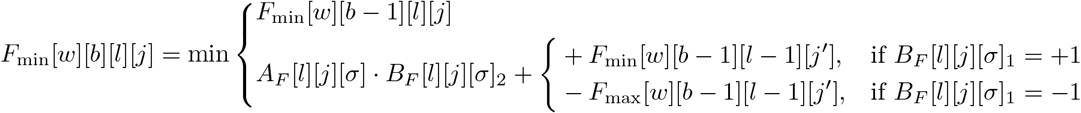

We first load all the values of *A_F_* [*l*][*j*][*σ*]*, B_F_* [*l*][*j*][*σ*]_2_,, 0 ≤ *j < d* into two registers. When iterating *j* from 0 to *d* − 1, the multiplication of *A_F_* [*l*][*j*][*σ*] and *B_F_* [*l*][*j*][*σ*]_2_ can be simplified as a single SIMD multiply instruction can directly get all the results. Besides, we can transform *B_F_* [*l*][*j*][*σ*]_1_ into single mask number(an integer) where each bit represents the selection of +*F*_min_[*w*][*b*−1][*l*−1][*j*′] or −*F*_max_[*w*][*b*− 1][*l* − 1][*j*′]. Similarly addition, maximum and minimum operations can also be replaced by SIMD instructions. In summary, transforming all operations in the recurrence into SIMD instructions reduces the time complexity of the entire iterating-steps to *O*(*Nnk*).

In the optimizing-step, calculating Ω*_i_*[*w*][*b*] can also benefit from similar SIMD instructions which results in *O*(1) time. Although Ψ_[_*w*][*b*] still requires *O*(*d*) running time, it is notably faster comparing to iterating-step and calculating Ω*_i_*[*w*][*b*]. Consequently, SIMD instructions contribute significantly to saving time in the overall seed calculation.

In detail, we require that *d* can not exceed 32 which allows to save at most 32 different 16-bits values in a 512-bits register. Regardless of the value of *d*, the overall running time remains nearly constant.

### 2.4 SubseqHash2r: Variant for Reverse Complement

In sequence analysis, it is a much desired property to *not* distinguish a sequence and its reverse complement. We can achieve this property by using symmetric tables in an ABCk order so that a length-*n* string and its reverse complement can be mapped into the same set of *k* seeds. We refer to this variant of SubseqHash2 as SubseqHash2r. Note that this symmetry is enabled by the multiple orders defined in the ABCk order, and hence cannot be directly transferred to SubseqHash.

Let ***x*** ∈ Σ*^n^* be a length-*n* string and let 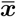 be its reverse complement. We aim for having that *i*-th seed of ***x*** to be equal to the (*k* − *i* + 1)-th seed of 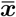, for every 1 ≤ *i* ≤ *k*, formally:

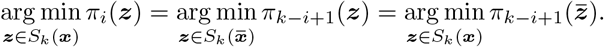

This condition is met if *π*_*i*_(***z***) = *π*_*k*−*i*+1_(***z***), for every 1 ≤ *i* ≤ *k*.

Here we derive the required property on the ABCk tables in order to generate the same set of seeds for a string and its reverse complement. Let ***z*** = *z*_1_*z*_2_ *… z_k_* and therefore 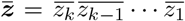, where we use 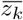 to represent the “complement” of letter *z_k_*. We require *π*_*i*_(***z***) = *π*_*k*−*i*+1_(***z***), for every 1 ≤ *i* ≤ *k*. Recall that 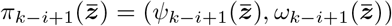 and

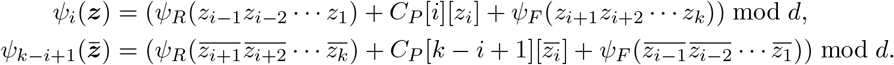

In order to make 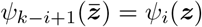 we require the following

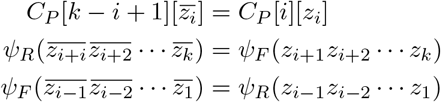

By the definitions of *ψ_F_* and *ψ_R_*, they lead to the following for every 1 ≤ *i* ≤ *k* and *z* ∈ Σ.

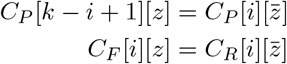

The same approach can be used to derive the requirements needed for making 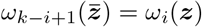, which we directly show below, for every 1 ≤ *i* ≤ *k, z* ∈ Σ, and 0 ≤ *j < d*:

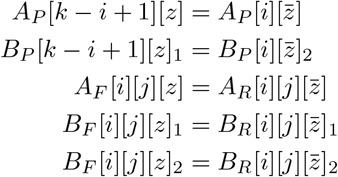

### 2.5 SubseqHash2w: Variant Concatenating a Substring

The recently developed method strobemer extracts a substring and some minimizers from the subsequent windows and concatenates them as a seed [23, 24, 14]. We apply this idea to SubseqHash2, creating a new variant termed SubseqHash2w. In SubseqHash2w, a seed is formed by two parts: a substring and the smallest subsequence from the following window. We use an additional parameter *k*_0_ to specify the length of the preceding substring. For an input ***x*** of length *k*_0_ + *n*, SubseqHash2w extracts the first substring of length *k*_0_, and then compute *k* seeds of length *k* from the remaining string of length *n* using the above algorithm. The leading substring of length *k*_0_ will be concatenated with each of the *k* seeds, resulting in *k* seeds of length *k*_0_ + *k*. In experiments, we compare SubseqHash2w with strobemer.

### 2.6 Selection of Parameters

SubseqHash2 has parameters *n, k, d*, and *t*. The combination of *n* and *k* balances the sensitivity and precision of the produced seeds. For a fixed *n*, a larger *k* provides lower sensitivity and fewer false positives. Different values of *n* may have different performance while larger *n* also takes more running time. The sensitivity can be increased if *d* is increased; with SubseqHash2 we can large *d* as its running time is independent of *d* for up to *d* = 32. Parameter *t* specifies the number of seeds in a window (length-*n* string), which should be in the range of [1*, k*]. The selection of *t* can be application-dependent; in general, data with higher error rates might need a larger *t* to produce enough seed-matches. The algorithm is implemented to allowing for specifying *t*, in which case the optimizing-step can be sped up to just generate *t* seeds. SubseqHash2w has an additional parameter *k*_0_, which also balances sensitivity and precision. In the experiments, we use *k*_0_ = *k/*2.

## 3 Results

We first present an analysis of the probability of hash collision as a function of edit distance for different seeding methods, illustrating their basic properties. Next, we benchmark the running time of SubseqHash and SubseqHash2. Last, we compare different seeding methods for three applications: read mapping, pairwise sequence alignment, and overlap detection.

### 3.1 Comparison of Probability of Hash Collision

Most seeding methods can be interpreted as a (hashing) function that maps a string (typically a window in a long sequence in practice) into a seed. For examples, minimizer maps a length-*n* string to its smallest length-*k* substring, and SubseqHash/SubseqHash2 maps a length-*k* string to its smallest length-*k* subsequence. It is desirable for such a seeding method *h* to be both sensitive and precise (aka locality-sensitive), i.e., the probability of hash collision Pr(*h*(***x***) = *h*(***y***)) is high (resp. low) if the edit distance between two strings ***x*** and ***y***, written as *edit*(***x***, ***y***), is small (resp. large). When repeating *h* by *t* times, a length-*n* string ***x*** will be mapped into *t* seeds, written as *h_i_*(***x***), 1 ≤ *i* ≤ *t*, and a seeding method is desired to achieve a high probability of having at least one hash collision Pr(∨_1≤*i*≤*t*_(*h_i_*(***x***) = *h_i_*(***y***)) when *edit*(***x***, ***y***) is small, and a high probability of not having any hash collision Pr(∧_1≤*i*≤*t*_(*h_i_*(***x***) ≠ *h_i_*(***y***)) when *edit*(***x***, ***y***) is large.

The probability of hash collisions can be estimated using simulations. We randomly generate pairs of strings over alphabet {A, C, G, T}. A simulated pair will be put into category-*i* if the edit distance between them is *i, i* = 1, 2*, …,* 10. We simulate 100,000 pairs in each category. We apply minimizer, SubseqHash, and SubseqHash2 on the simulated pairs with repetition (*t* = 10 or *t* = *k*) and without repetition (*t* = 1). Minimizer is repeated by using different random seeds in its hash function in defining the order for kmers. SubseqHash is repeated by using random, independent tables used in the ABC order. SubseqHash2 generates *t* sets of seeds in a single run. The frequency of hash collisions (at least one hash collision in the case of repetitions) in each category will be used to estimate the probability.

The estimated probabilities are shown in Figure 2 and Supplementary Figures 1, 2, 3. First, observe that SubseqHash2 and SubseqHash with the same number of repeats exhibits nearly identical probability. This is exactly the goal of SubseqHash2: the same performance as SubseqHash but running in a much reduced time complexity. Second, repeating is effective for SubseqHash2 and SubseqHash, demonstrated by the facts that the probability can be drastically increased with small edit distance, while remaining nearly zero with large edit distance. On the contrary, repeating is not effective for minimizer, as the probability remains low even with small edit distance. We also included the results of using all kmers as seeds (therefore a pair of string has at least one hash collision with “all kmer” if they share at least one kmer). As expected, the probability of repeating minimizers is close to and bounded by that of using all kmers. For experiments in the following Sections, we simply include the results of “all kmers” to approximate the results of repeating minimizer.

**Figure 2.**
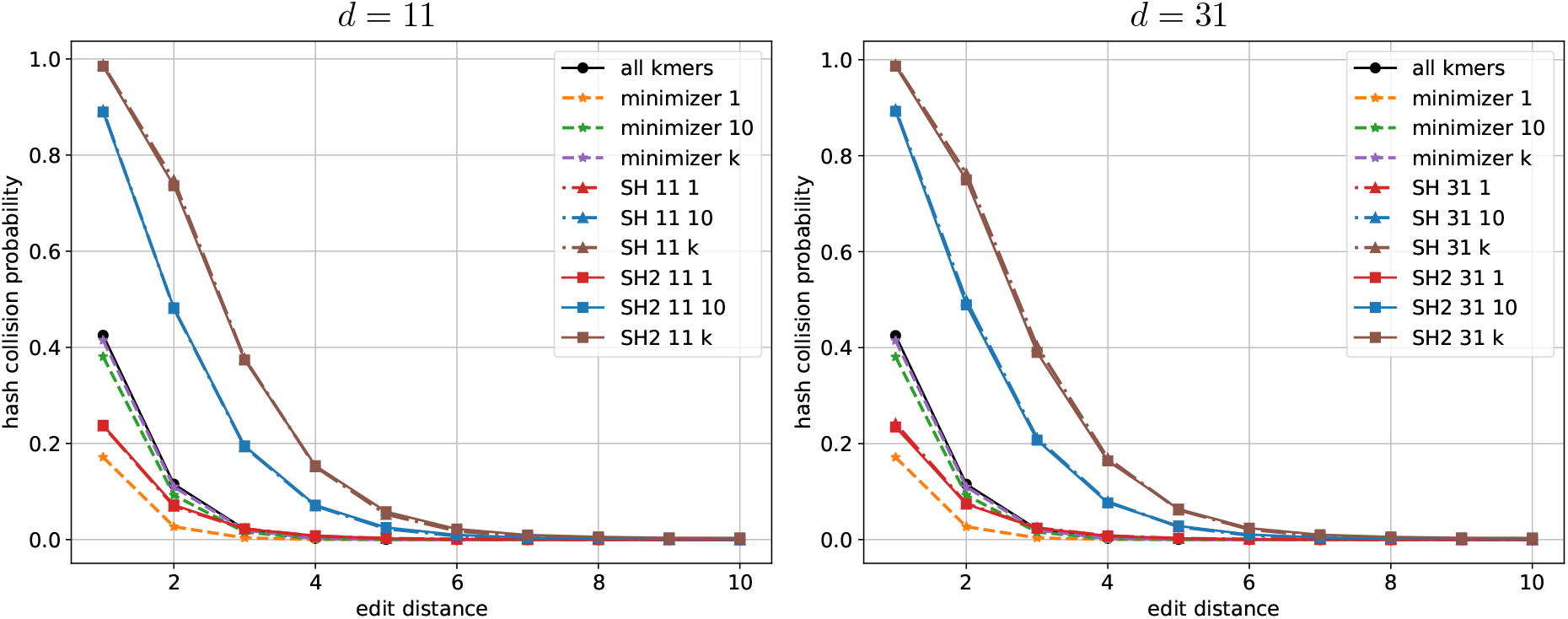
The probability of hash collision (at least one hash collision in the case of *t* = 10 and *t* = *k* = 24) estimated for different seeding methods with *n* = 30 and *k* = 24. “minimizer *t*” represents repeating minimizer *t* times. “SH *d t*” and “SH2 *d t*” represent SubseqHash and SubseqHash2 with parameter *d* and repeating *t* times.

**Figure 3.**
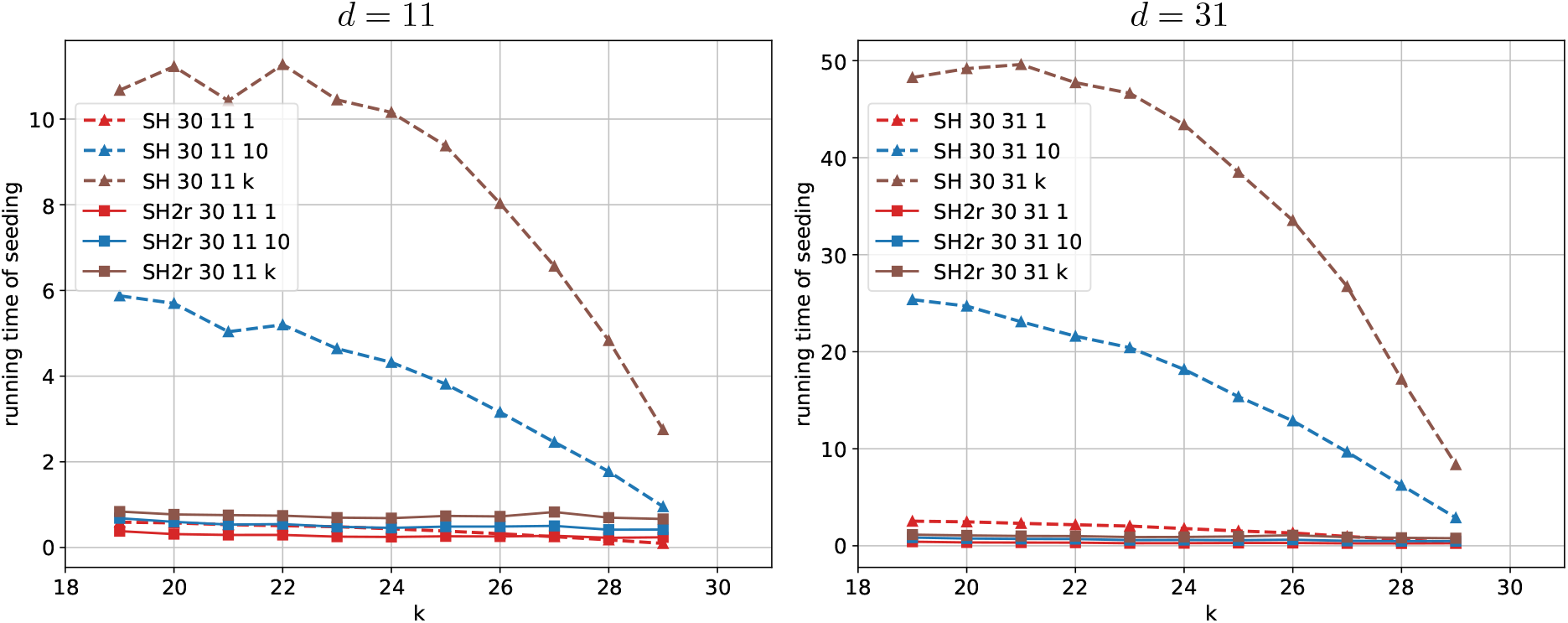
The average CPU time (second) of SubseqHash and SubseqHash2 per read. “SH *n d t*” and “SH2r *n d t*” represent SubseqHash and SubseqHash2r with window size of *n*, parameter *d*, and repeating *t* times; for both cases the lines connect points with varying *k*.

**Figure 4.**
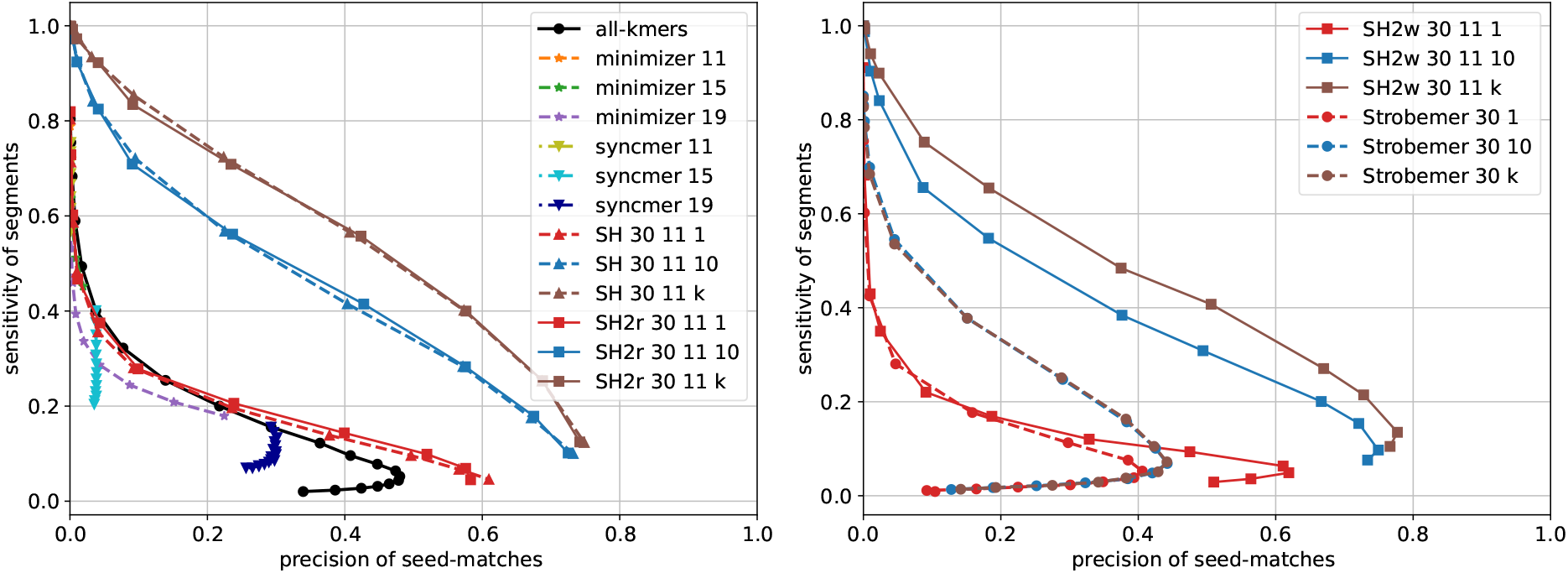
Comparison of averaged precision of seed-matches and sensitivity of segments. In the left figure, line “all-kmers” connects points with varying *k* from 10 to 25; “minimizer *n*” represents a window size of *n*, with points on it corresponding to varying *k*. “syncmer *k*” represents closed syncmer with a length-*k* seed; the line connects points with varying *s* from 2 to *k* − 1 In the right figure, “SH2w *n d t*” represents SubseqHash2w with window size of *n*, parameter *d*, and repeating *t* times; *k*_0_ = *k/*2. “Strobemer *n t*” represents Strobemer (two windows) with window size of *n* and repeating *t* times; the line connects points with varying parameter *k* from 5 to 20.

**Figure 5.**
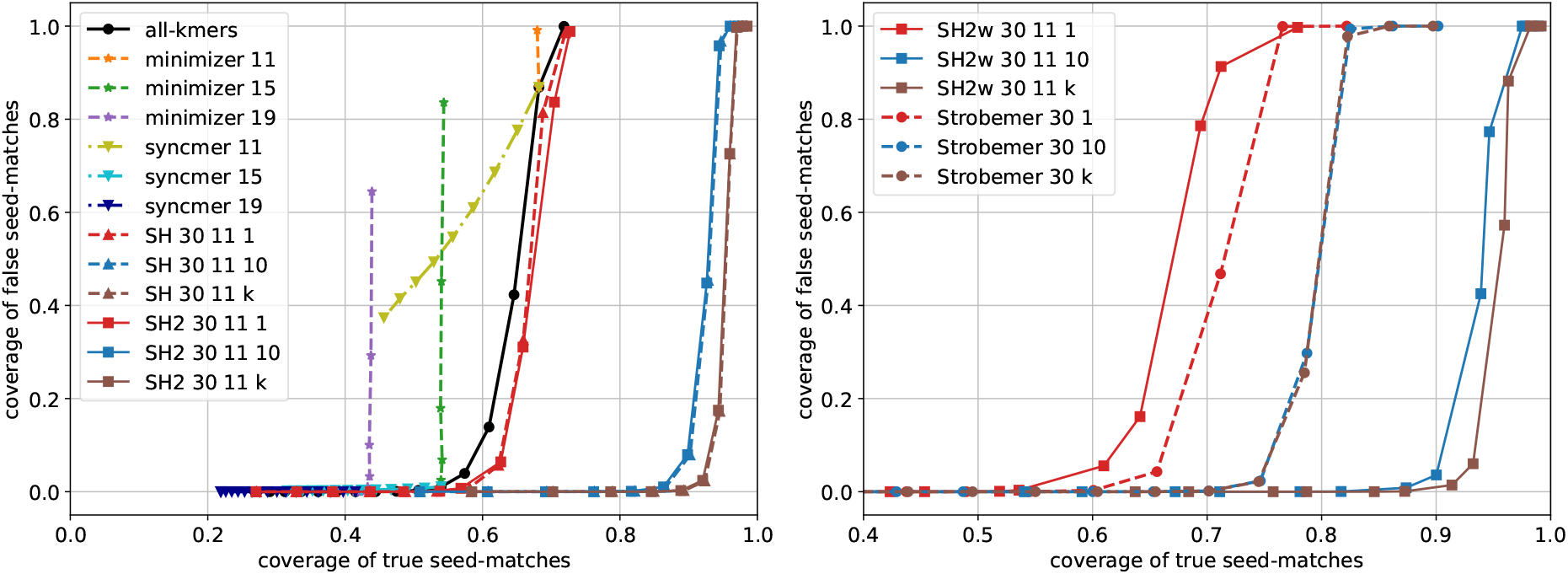
The true and false coverages of different seeding methods on data with error rate *r* = 10%. The same parameters as in Section 3.3 are used.

**Figure 6.**
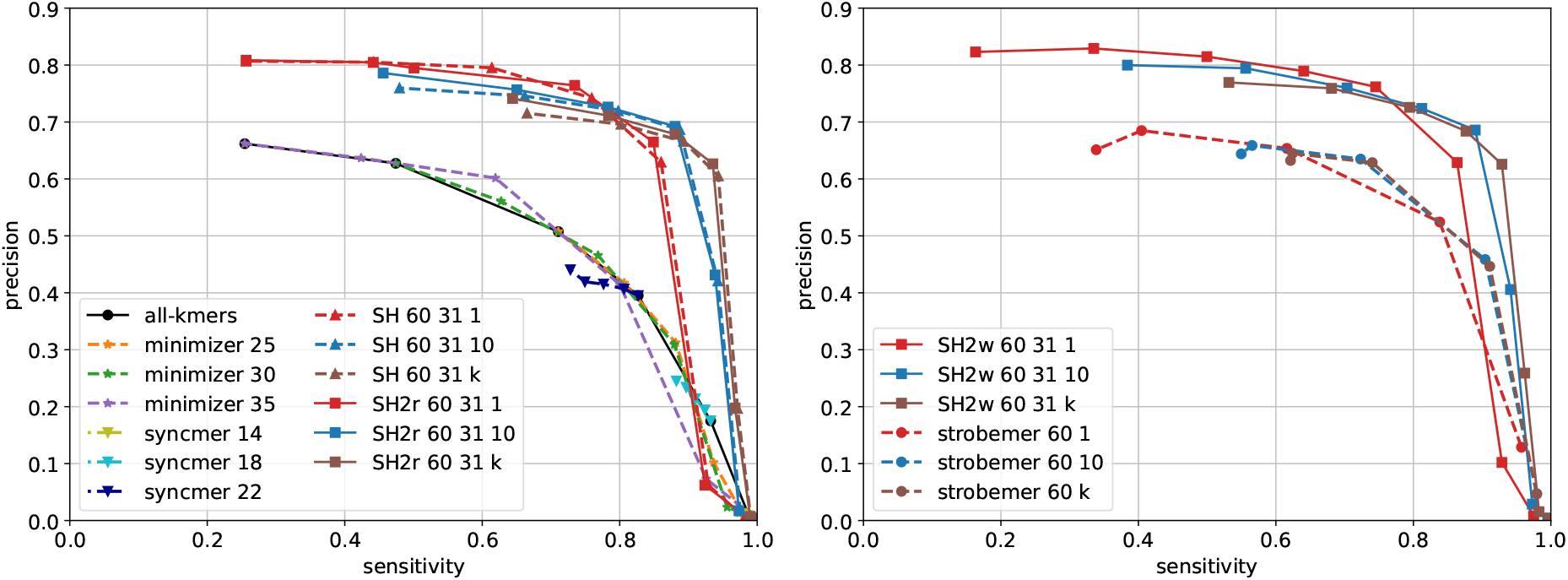
Overlap detection results on 10,000 reads sampled from the *E. coli* SRX533603 dataset with window size 60 and *d* = 31. For minimizers with window sizes *n* = 25, 30, 35, each curve is drawn with six seed lengths *k* evenly spaced between 10 and *n* (when *k* = *n*, all *k*-mers are in the seed set). For syncmers with seed lengths *k* = 14, 18, 22, each curve is draw with five *s*-mer lengths evenly spaced between 2 and *k* (when *s* = *k*, all *k*-mers are selected). For SubseqHash, SubseqHash2r, strobemer, and SubseqHash2w, the window size is *n* = 60. For SubseqHash, SubseqHash2r, and SubseqHash2w, the seed lengths *k* = ⌊*rn*⌋ for *r* = 0.5, 0.55, 0.6, 0.65, 0.7, 0.75, 0.8, 0.85 are plotted. For strobemer, a seed consists of two *k*-mers for *k* = ⌊*rn*/2⌋ with *r* = 0.35, 0.45, 0.55, 0.65, 0.75, where the first *k*-mer is the leading *k*-mer in the window, and the second *k*-mer is selected by the randstrobe algorithm from the remaining part of the window. For SubseqHash2w, a seed of length *k* consists of the leading *k*_0_ = ⌊*k*/3⌋-mer in the window and a SubseqHash2 seed of length ⌊2*k*/3⌋ from the remaining part of the window.

### 3.2 Comparison of Running Time

We compare the running time of SubseqHash and SubseqHash2 in Figure 3 and Supplementary Figure 4. Both methods are applied to the SRX499318 dataset (with PacBio long reads), and we report the average CPU time per read. With the same parameters *d* and *t*, SubseqHash2 can be 50 times faster than SubseqHash. SubseqHash2 repeating *k* times can be even faster than SubseqHash without repeating. It is also noteworthy that the running time of SubseqHash2 only experiences a slight increase as *d* and *t* grow, suggesting that the efficiency of SubseqHash2 remains nearly independent of *d* and *t*, confirming our theoretical analysis that the time complexity is substantially improved.

We also compare with kmer-based methods on the same dataset. Not surprisingly, kmers, Minimizers, and syncmers are exceedingly fast, taking less than 5 seconds across almost all choices of parameters, as they only need a linear scan of each read. Strobemers at a typical configuration (two strobes of length 15 within a window of size 50) consumes 145.19 seconds with 10 repeats. In comparison, SubseqHash2, configured with *n* = 30, *k* = 25, *d* = 31, and 10 repeats, uses 577.45 seconds, placing it in a comparable time range as Strobemers. Notably, SubseqHash, with the identical (*n, k, d, t*) as SubseqHash2, requires a substantially longer running time of 15,364 seconds to complete the task. We note that repeat is a sensible option for both Strobemers and SubseqHash2, in order to achieve their best performances in three following applications.

### 3.3 Read Mapping

we now consider the task of mapping error-prone long reads to reference genomes. For this purpose, we simulate a set of reads using PBSIM2 [20] from chromosome X of Drosophila melanogaster (AE014298.5) that mirrors the sequencing error profile (reported in [27]) of the PacBio SRX499318 dataset. Reads that share long substrings with the parts of the reference where they originate can be mapped easily using either seeding methods. The real challenge lies in identifying true seed matches for reads (or parts of reads) that barely share any meaningfully long substring with the reference. Our seeding method is designed to tackle such “hard” reads. We argue that this is a valid and intended pipeline—it would be unwise to completely abandon the agile substring-based methods with existing highly optimized implementations such as minimap2 [11] for the easier tasks; instead, we use them as a filter to expose cases where only subsequence-based seeds can provide a satisfactory solution. From all the simulated reads, 4, 295 unmapped read fragments from the results of minimap2 that are longer than 1000 bp are kept for our analysis.

SubseqHash2 is compared with minimizer, closed syncmer, and all-kmers, for generating seeds on both the reads and the reference chromosome. As we do not know the strand of reads, we use SubseqHash2r, with one run being able to cover both strands; ther methods need to be applied both on the reads and their reverse complements. All seeding methods are set to generate seeds of length *k* from sliding windows of length *n*; seed-matches are then collected between the reads and the reference. When repeats are used, seed-matches from repeated runs are pooled. Note that a seed-match specifies *k* aligned characters across the two sequences. We define a seed-match to be *true* if over 50% of its *k* aligned characters coincide with the ground truth;otherwise, we call it a false seed-match.

In the seed-extend or seed-chain-extend framework of read mapping, more true seed-matches and fewer false seed-matches are definitely desirable. We therefore report the *precision of seed-matches*, defined as the number of true seed-matches divided by the total number of seed-matches. Additionally, it is also crucial that these seed-matches can be evenly distributed across the read, ensuring sufficient information for chaining and producing a complete alignment. To assess this, we partition all reads into segments of length 200 bp. An ideal seeding would cover every segment with true seed-matches. We hence also report a *segment sensitivity*, defined as the percentage of segments with at least one true seed-match.

In Figure 4 and Supplementary Figure 5, we present these metrics acquired by various methods. Observe that the segment sensitivity of substring-based methods drops rapidly as *k* increases. This is because in these “hard” reads the exact match of long kmers with reference is infrequent, as otherwise minimap2 would align them correctly. In contrast, SubseqHash2 with repeats demonstrates the ability to capture adequate true seed matches while precision increases, resulting in a curve significantly above others. Furthermore, the curves of substring-based methods confirm that on “hard” regions, no choice of *k* can produce a satisfactory balance between sensitivity and precision; whereas SubseqHash2 with repeats achieves a four times higher sensitivity at the peak precision of kmers. This attests to the superiority of SubseqHash2 in generating high-quality seeds for aligning difficult reads.

We compare SubseqHash2w with Strobemer (Randstrobe [23] with two windows), as both employ multiple windows. To repeat Strobemer, we change the random seed in its hash function. Without repeating SubseqHash2w shows slightly better performance. However, with repetitions, SubseqHash2w demonstrates a substantial improvement, highlighting the effectiveness of repetition in SubseqHash and SubseqHash2.

### 3.4 Pairwise Sequence Alignment

We then evaluate the performance of seeding methods for pairwise sequence alignment through simulations. For each error rate *r* = 0.05, 0.1, 0.15, we simulate 10 pairs of long sequences and report the average measures. We simulate a pair by randomly generating a sequence of length *L* = 1, 000, 000 as its first sequence, followed by applying an edit, with probability of *r/*3 being a substitution, insertion, or deletion, at each position, to get its second sequence. We compute the *coverage* of true (resp. false) seed-matches, defined as the proportion of characters encompassed by true (resp. false) seed-matches. High true coverage and low false coverage are desirable, as the former implies long-range of correct alignment while the latter implies low chance of producing incorrect alignment. These two measures can be interpreted as sensitivity and precision of a seeding method for sequence alignment.

We compare SubseqHash2 with other four methods: minimizers, closed syncmer, all-kmers, and SubseqHash. The results are given in Figure 5 and Supplementary Figures 6, 7, 8. Again, SubseqHash2 mirrors the accuracy from SubseqHash at the same level of repetitions, as we aimed for. With repetitions, SubseqHash2/SubseqHash considerably outperforms substring-based methods at all error rates. The results of SubseqHash2w and Strobemer are also shown. Although Strobemer performs better without repeating, SubseqHash2w significantly outperforms with repetitions.

### 3.5 Overlap Detection

Overlap graph is the fundamental data structure used by many long-read genome assemblers where each pair of overlapping reads are connected. Accurately identifying such pairs is thus crucial: having too many false positives tends to produce a tangled graph, while missing true positives can cause gaps in the assembly. Seeding methods are used to alleviate the computational burden by efficiently filtering out a large fraction of non-overlapping pairs: each read is first transformed into a set of seeds; pairs of reads are considered candidates for further verification only if they share a common seed. Usually, heuristics are applied such as filtering out high-frequency seeds, requiring more seed-matches to trigger a candidate pair, chaining co-linear seed-matches, and applying other fine-grained verification methods (e.g., local alignment), all of which can improve the final overlap results. We note that such pre- and post-processing steps are independent of the choice of seeds, so we omit them in this experiment to have a direct comparison of performances among different seeding schemes.

Two PacBio long reads datasets, SRX533603 (*E. coli*) and SRX499318 (*D. melanogaster*) from [4] are used. Reads are first mapped to reference genomes with minimap2 to construct a ground-truth for evaluation. From both datasets, 10, 000 reads are sampled from those that are confidently mapped. A pair of reads are considered truly overlap (i.e., ground-truth) if their mapped regions on the reference overlap by at least 15bp. Sensitivity and precision are defined as the fractions of reported pairs that are correct over all ground-truth pairs and over all reported pairs, respectively. Again, in order to be able to identify overlapping reads from opposite strands, with the exception of SubseqHash2r, other methods collect seeds twice from each read, one for each strand.

The precision-sensitivity curves are compared in Figure 6 and Supplementary Figures 9–11. SubseqHash and SubseqHash2 provide significantly higher sensitivity than substring-based seeds at the same level of precision, except when precision approaches zero where all methods achieve almost perfect sensitivity at the cost of reporting an excessive amount of false-positive pairs, which defeats the purpose of seeding. For substring-based seeds, the extreme of repeating is to include all-kmers, which produces little improvement. Repeating helps (random) Strobemers but only to a limited extent (observe that strobemer curves with 10 and *k* repeats almost overlap). This is because its components are kmers and hence have a restricted range of choices; furthermore, all the kmers in two similar windows can be easily destroyed by just a few errors, in which case no matching Strobemers can be found regardless of the number of repeats. In contrast, the performance of SubseqHash/SubseqHash2 can still benefit from more repeats, justifying the motivation of producing *k* seeds in one run.

## 4 Discussion

We introduced SubseqHash2, an advanced subsequence-based seeding method. Remarkably, it can be 50 times faster than its predecessor, SubseqHash, while preserving high seed quality. The innovative strategy of using a pivot position allows SubseqHash2 to produce up to *k* seeds for a window in a single pass. This approach substantially reduces the computational time of producing *t* ≤ *k* sets of seeds from *O*(*Nnkdt*) to *O*(*Nnkd*) and grants SubseqHash2 the flexibility to adapt its seed count to enhance performance in various sequence analysis tasks without incurring additional computation. Additionally, we employ SIMD instructions in the dynamic programming algorithm for seed computation, enabling parallel computation of *d* subproblems and further reducing the dynamic programming running time to *O*(*Nnk*). Throughout our experiments, SubseqHash2 matches the performance of SubseqHash while delivering a significant reduction in actual running time. Moreover, these algorithmic innovations allow for generating the same sets of seeds for a sequence and its reverse complement, which further halves the running time for reads with unknown strand information. SubseqHash2 is versatile, can be combined with kmers in seeding as needed.

We recognize that the superior seed quality of our approach comes at the cost of a more complicated algorithm, and therefore, despite being substantially improved, still slower comparing to the simple and fast substring-based seeding methods. We are actively exploring ideas for further acceleration, both in terms of algorithmic design and implementation optimization. On the other hand, we believe strategically combining the agility of substring-based methods and the high sensitivity of subsequence-based seeding can potentially produce an ideal solution for analyzing data with high mutation/error rates, as demonstrated by our read mapping experiment. Given the unsatisfied accuracy of substring-based methods on error-prone data, as well as the common struggle of choosing *k*, we are optimistic that SubseqHash2, as a less swift but more accurate alternative, can establish its value across a broader spectrum of applications.

## Acknowledgment

This work is supported by the US National Science Foundation (2019797 and 2145171 to M.S.) and by the US National Institutes of Health (R01HG011065 to M.S.).

## A Algorithm for calculating Ψ*_i_*[*w*][*b*]

Calculating Ψ*_i_*[*w*][*b*] goes down to solve this abstracted problem: given integer *d*, integer *d*_0_, and two sorted lists *V*_1_ and *V*_2_ for which all elements are in {0, 1*, …, d* − 1}, calculate 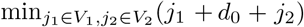 mod *d*. The constant *d*_0_ stands for *C_P_* [*i*][*X*_*w*+*b*+1_] in the original task. Below we design two algorithms: the first one runs in *O*(*d*) time assuming that a 2*d* bit-vector can be fit in a machine word; the second algorithm runs in *O*(*d ·* log *d*) solves this problem for arbitrarily large *d*.

**Algorithm 1.** We first transfer *V*_1_ into a number *x*_1_ stored as a *d*-bit vector:

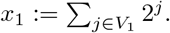

We use *x*_1_[*j*] to represent the *j*-th bit of its underlying bit-vector, formally defined as *x*_1_[*j*] := *lx*_1_*/*2*^j^J* mod 2. Clearly, for any 0 ≤ *j < d*, we have *x*_1_[*j*] = 1 if and only if *j* ∈ *V*_1_.

We then calculate *x*, defined below. For each element *j* ∈ *V*_2_ we shift the bit-vector of *x*_1_ to the left by *j* bits to get a new bit-vector (of size 2*d* bits, with possible padding 0s to its left). We calculate *x* by bit-wise OR over these new bit-vectors. Clearly, *x* is of size 2*d* bits.

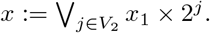

We now show that, the *i*-th bit of *x* equals to 1, i.e., *x*[*i*] = 1, if and only if there exists *j*_1_ ∈ *V*_1_ and *j*_2_ ∈ *V*_2_, such that *j*_1_ + *j*_2_ = *i*. Considering how *x* is calculated, we have that *x*[*i*] = 1 if and only if there must be *j*_2_ ∈ *V*_2_ such that 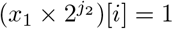. We can write:

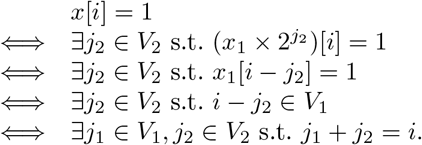

Now we transform *x* back to a set *V* by defining *x*[*i*] = 1 if and only if *i* ∈ *V*. In other words, *V* now stores all the possible values that can be obtained by *j*_1_ + *j*_2_ where *j*_1_ ∈ *V*_1_ and *j*_2_ ∈ *V*_2_. The final solution can be calculated by enumerating elements in *V* :

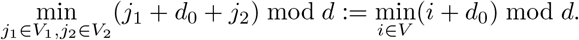

In this algorithm, computing *x*_1_ takes *O*(*d*) time; calculating *x* takes *O*(*d*) time as well because there are at most *d* multiplication (bit-shifting) and *d* bitwise OR operations, which of which can be done in *O*(1) time. The final minimization takes *O*(*d*) time. Therefore, the overall running time of this algorithm is *O*(*d*).

**Algorithm 2.** We have another *O*(*d ·* log *d*) algorithm for general cases. For each value *j*_1_ in *V*_1_, the optimal value from *V*_2_ is *j*\* := (2*d* − *j*_1_ − *d*_0_) mod *d*, as in this case (*j*_1_ + *j*\* + *d*_0_) mod *d* = 0. But *j*\* might not exist in *V*_2_. To find the optimal value in *V*_2_, we note that for each *j*_2_ ∈ *V*_2_ the objective function can be represented in a simple closed form:

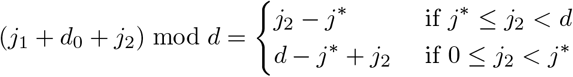

This closed form can also be illustrated and verified using the table given below.

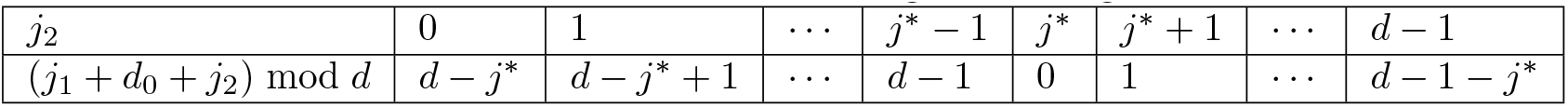

For each *j*_1_, above closed form gives an easy way to find optimal value in *V*_2_: among these *j*_2_ in *V*_2_ satisfying *j*\* ≤ *j*_2_, the optimal one (which gives the smallest objective value) would be the smallest such *j*_2_; among these *j*_2_ in *V*_2_ satisfying *j*_2_ *< j*\*, the optimal one (which gives the smallest objective value) would be the smallest such *j*_2_ as well. Note that, for any *j*_2_ ≥ *j*\*, it always gives a smaller objective value than any *j*_2_ *< j*\*.

Hence, we can apply binary search on *V*_2_ to find the optimal *j*_2_. Specifically, we search *V*_2_ to find the first value that is equal to or larger than *j*\*. If it exists, denoting it as *p*, we calculate the corresponding objective value as *p* − *j*\*. Otherwise, we find the smallest value in *V*_2_ (which can be done in *O*(1) time as we assume both *V*_1_ and *V*_2_ are sorted), denoting it as *q*, we calculate the corresponding objective value as *d* − *j*\* + *q*. The overall optimal objective value will be the smallest one by enumerating all *j*_1_ ∈ *V*_1_.

For each *j*_1_ ∈ *V*_1_, above binary search takes *O*(log *d*) time; the entire algorithm runs in *O*(*d ·* log *d*) time.

## B An example of the ABCk order

We give an example for the ABCk order, determined by the random tables shown in Tables 1, 2, and 3 with *k* = 6, *d* = 5, and Σ = {*A, C, G, T* }. The score of ***z*** = CTAACT can be calculated following the recurrences, with detailed given below. We first compute *ψ*_*F*_, *ω*_*F*_ using tables *A*_*F*_, *B*_*F*_, and *C*_*F*_, and *ψ*_*R*_, *ω*_*R*_ with tables *A*_*R*_, *B*_*R*_, and *C*_*R*_:

| | Reverse | Pivot | Forward | $\psi_R$ | $\omega_R$ | $\psi_F$ | $\omega_F$ |
| --- | --- | --- | --- | --- | --- | --- | --- |
| $\pi_1$ | ∅ | C | TAACT | 0 | 0 | 3 | -155 |
| $\pi_2$ | C | T | AACT | 3 | -18 | 3 | -52 |
| $\pi_3$ | TC | A | ACT | 0 | -37 | 2 | -156 |
| $\pi_4$ | ATC | A | CT | 4 | 1 | 2 | -36 |
| $\pi_5$ | AATC | C | T | 0 | 9 | 0 | 79 |
| $\pi_6$ | CAATC | T | ∅ | 2 | -191 | 0 | 0 |

**Table 1.** An example of random tables *A*_*P*_, *B*_*P*_, *C*_*P*_ used in an ABCk order. Entries in table *A*_*P*_ are integers drawn from [−100, −10] ∪ [10, 100].

| $i$ | $\sigma$ | | | |
| --- | --- | --- | --- | --- |
|  | A | C | G | T |
| 1 | -24 | 82 | -34 | 11 |
| 2 | -90 | 85 | 33 | -95 |
| 3 | -12 | 89 | 29 | -12 |
| 4 | 49 | 71 | -73 | -18 |
| 5 | -84 | -36 | 91 | 70 |
| 6 | 49 | -17 | -32 | 16 |

**(b)** Table $B_P$
| $i$ | $\sigma$ | | | |
| --- | --- | --- | --- | --- |
|  | A | C | G | T |
| 1 | (-1, 1) | (-1, -1) | (1, 1) | (1, -1) |
| 2 | (-1, 1) | (-1, -1) | (1, -1) | (1, 1) |
| 3 | (-1, 1) | (1, -1) | (-1, -1) | (1, 1) |
| 4 | (-1, -1) | (1, 1) | (-1, 1) | (1, -1) |
| 5 | (1, 1) | (1, -1) | (-1, 1) | (-1, -1) |
| 6 | (1, -1) | (-1, -1) | (1, 1) | (-1, 1) |

**(c)** Table $C_P$
| $i$ | $\sigma$ | | | |
| --- | --- | --- | --- | --- |
|  | A | C | G | T |
| 1 | 3 | 1 | 4 | 2 |
| 2 | 3 | 1 | 4 | 0 |
| 3 | 3 | 1 | 4 | 0 |
| 4 | 0 | 2 | 4 | 3 |
| 5 | 3 | 1 | 4 | 2 |
| 6 | 4 | 3 | 0 | 2 |

**Table 2.**
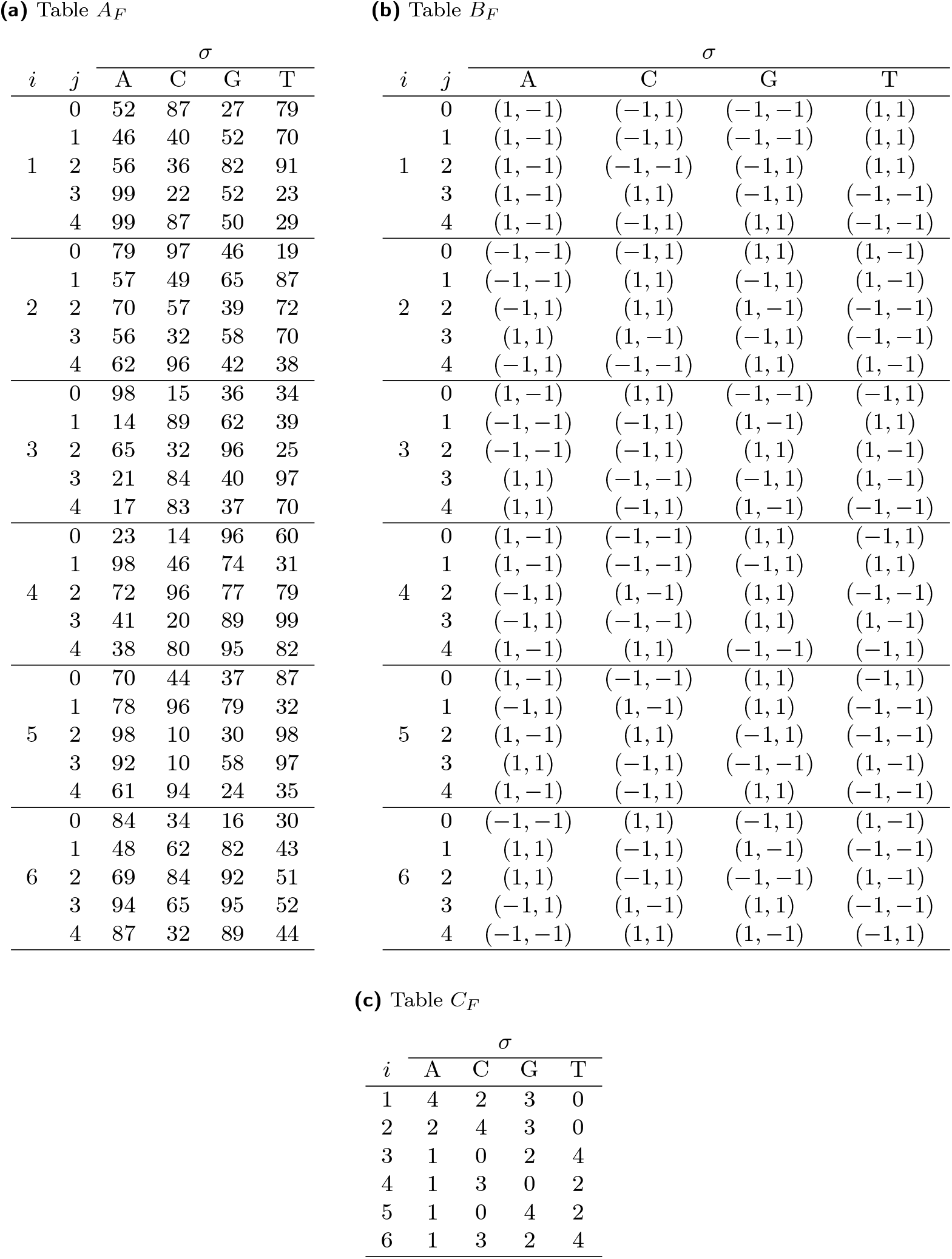
An example of random tables *A*_*F*_, *B*_*F*_, *C*_*F*_ used in an ABCk order. Entries in table *A_F_* are integers drawn from [10, 100].

**Table 3.** An example of random tables *A*_*R*_, *B*_*R*_, *C*_*R*_ used in an ABCk order. Entries in table *A*_*R*_ are integers drawn from [10, 100].

| $i$ | $j$ | $\sigma$ | | | |
| --- | --- | --- | --- | --- | --- |
|  |  | A | C | G | T |
| 1 | 0 | 15 | 44 | 68 | 42 |
|  | 1 | 28 | 47 | 71 | 79 |
|  | 2 | 48 | 48 | 88 | 14 |
|  | 3 | 64 | 18 | 99 | 60 |
|  | 4 | 11 | 92 | 74 | 96 |
| 2 | 0 | 15 | 23 | 92 | 33 |
|  | 1 | 96 | 35 | 89 | 87 |
|  | 2 | 12 | 67 | 21 | 83 |
|  | 3 | 43 | 29 | 85 | 99 |
|  | 4 | 39 | 18 | 14 | 70 |
| 3 | 0 | 37 | 48 | 60 | 99 |
|  | 1 | 84 | 24 | 98 | 19 |
|  | 2 | 51 | 23 | 35 | 79 |
|  | 3 | 66 | 91 | 91 | 26 |
|  | 4 | 29 | 67 | 93 | 31 |
| 4 | 0 | 46 | 46 | 91 | 94 |
|  | 1 | 14 | 49 | 73 | 31 |
|  | 2 | 49 | 94 | 16 | 44 |
|  | 3 | 48 | 71 | 66 | 62 |
|  | 4 | 35 | 12 | 67 | 35 |
| 5 | 0 | 93 | 38 | 65 | 86 |
|  | 1 | 96 | 66 | 91 | 80 |
|  | 2 | 91 | 42 | 33 | 72 |
|  | 3 | 11 | 41 | 45 | 45 |
|  | 4 | 60 | 23 | 29 | 24 |
| 6 | 0 | 79 | 95 | 17 | 44 |
|  | 1 | 26 | 63 | 65 | 23 |
|  | 2 | 77 | 36 | 71 | 48 |
|  | 3 | 29 | 91 | 49 | 63 |
|  | 4 | 99 | 31 | 92 | 71 |

**(b)** Table $B_R$
| $i$ | $j$ | $\sigma$ | | | |
| --- | --- | --- | --- | --- | --- |
|  |  | A | C | G | T |
| 1 | 0 | (-1, -1) | (1, -1) | (-1, 1) | (1, 1) |
|  | 1 | (-1, -1) | (1, -1) | (1, 1) | (-1, 1) |
|  | 2 | (-1, -1) | (1, -1) | (1, 1) | (-1, 1) |
|  | 3 | (-1, -1) | (1, -1) | (1, 1) | (-1, 1) |
|  | 4 | (-1, -1) | (1, -1) | (-1, 1) | (1, 1) |
| 2 | 0 | (1, -1) | (-1, -1) | (-1, 1) | (1, 1) |
|  | 1 | (1, 1) | (-1, -1) | (1, -1) | (-1, 1) |
|  | 2 | (1, -1) | (-1, 1) | (-1, -1) | (1, 1) |
|  | 3 | (-1, -1) | (1, 1) | (-1, 1) | (1, -1) |
|  | 4 | (-1, -1) | (1, -1) | (1, 1) | (-1, 1) |
| 3 | 0 | (1, 1) | (1, -1) | (-1, -1) | (-1, 1) |
|  | 1 | (1, -1) | (-1, -1) | (1, 1) | (-1, 1) |
|  | 2 | (-1, 1) | (1, 1) | (-1, -1) | (1, -1) |
|  | 3 | (1, -1) | (1, 1) | (-1, -1) | (-1, 1) |
|  | 4 | (-1, -1) | (1, -1) | (1, 1) | (-1, 1) |
| 4 | 0 | (-1, -1) | (1, -1) | (-1, 1) | (1, 1) |
|  | 1 | (1, 1) | (-1, 1) | (1, -1) | (-1, -1) |
|  | 2 | (1, 1) | (-1, 1) | (-1, -1) | (1, -1) |
|  | 3 | (1, 1) | (1, -1) | (-1, -1) | (-1, 1) |
|  | 4 | (1, 1) | (-1, 1) | (-1, -1) | (1, -1) |
| 5 | 0 | (-1, -1) | (-1, 1) | (1, 1) | (1, -1) |
|  | 1 | (1, 1) | (-1, 1) | (1, -1) | (-1, -1) |
|  | 2 | (-1, 1) | (1, -1) | (1, 1) | (-1, -1) |
|  | 3 | (-1, -1) | (1, 1) | (1, -1) | (-1, 1) |
|  | 4 | (-1, -1) | (1, -1) | (1, 1) | (-1, 1) |
| 6 | 0 | (-1, 1) | (-1, -1) | (1, 1) | (1, -1) |
|  | 1 | (-1, 1) | (-1, -1) | (1, 1) | (1, -1) |
|  | 2 | (-1, -1) | (1, 1) | (-1, 1) | (1, -1) |
|  | 3 | (1, -1) | (-1, -1) | (1, 1) | (-1, 1) |
|  | 4 | (1, 1) | (-1, 1) | (1, -1) | (-1, -1) |

**(c)** Table $C_R$
| $i$ | $\sigma$ | | | |
| --- | --- | --- | --- | --- |
|  | A | C | G | T |
| 1 | 0 | 3 | 1 | 2 |
| 2 | 4 | 3 | 1 | 2 |
| 3 | 4 | 2 | 1 | 0 |
| 4 | 2 | 1 | 4 | 3 |
| 5 | 0 | 3 | 4 | 2 |
| 6 | 3 | 1 | 0 | 2 |

Then we can calculate the values of *k* score functions *π*_1_, *π*_2_, *…*, *π*_6_ for ***z*** = CTAACT based on *ψ_F_*, *ω_F_*, *ψ_R_, ω_R_*, and table *A_P_*, *B_P_*, *C_P_* .

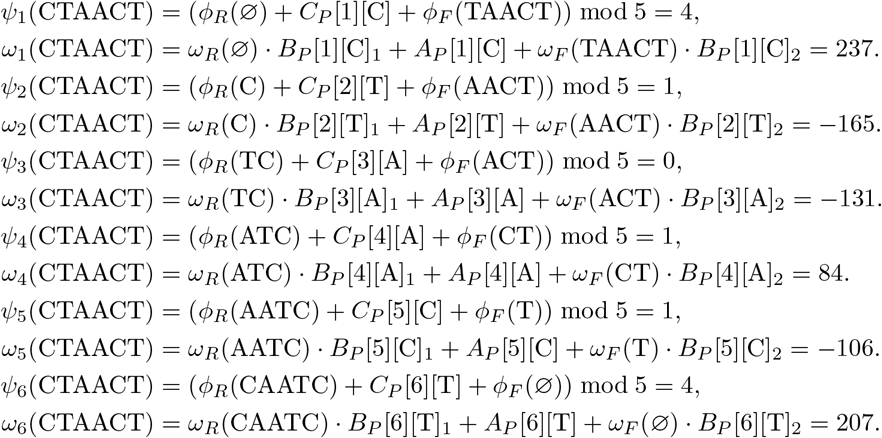

The *k* scores of ***z*** = CTAACT are listed below. Similarly, we also get the scores of ***z*′** = CCAACT.

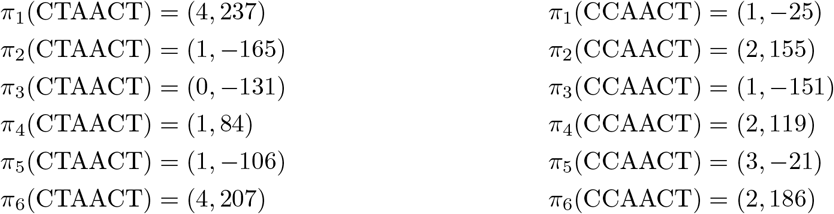

Note that the edit distance between ***z*** and ***z*′** is 1, but all the *k* scores are drastically changed. This is a desired property and is the goal of the design of the ABCk orders. Because of this, each of the *k* orders in the ABCk order behaves similarly to a pure random order, so that the probability of hash collision between two strings can be approximately the Jaccard index between the two sets of length-*k* subsequences.

