## Supplementary Figures for "Efficient Seeding for Error-Prone Sequences with SubseqHash2"

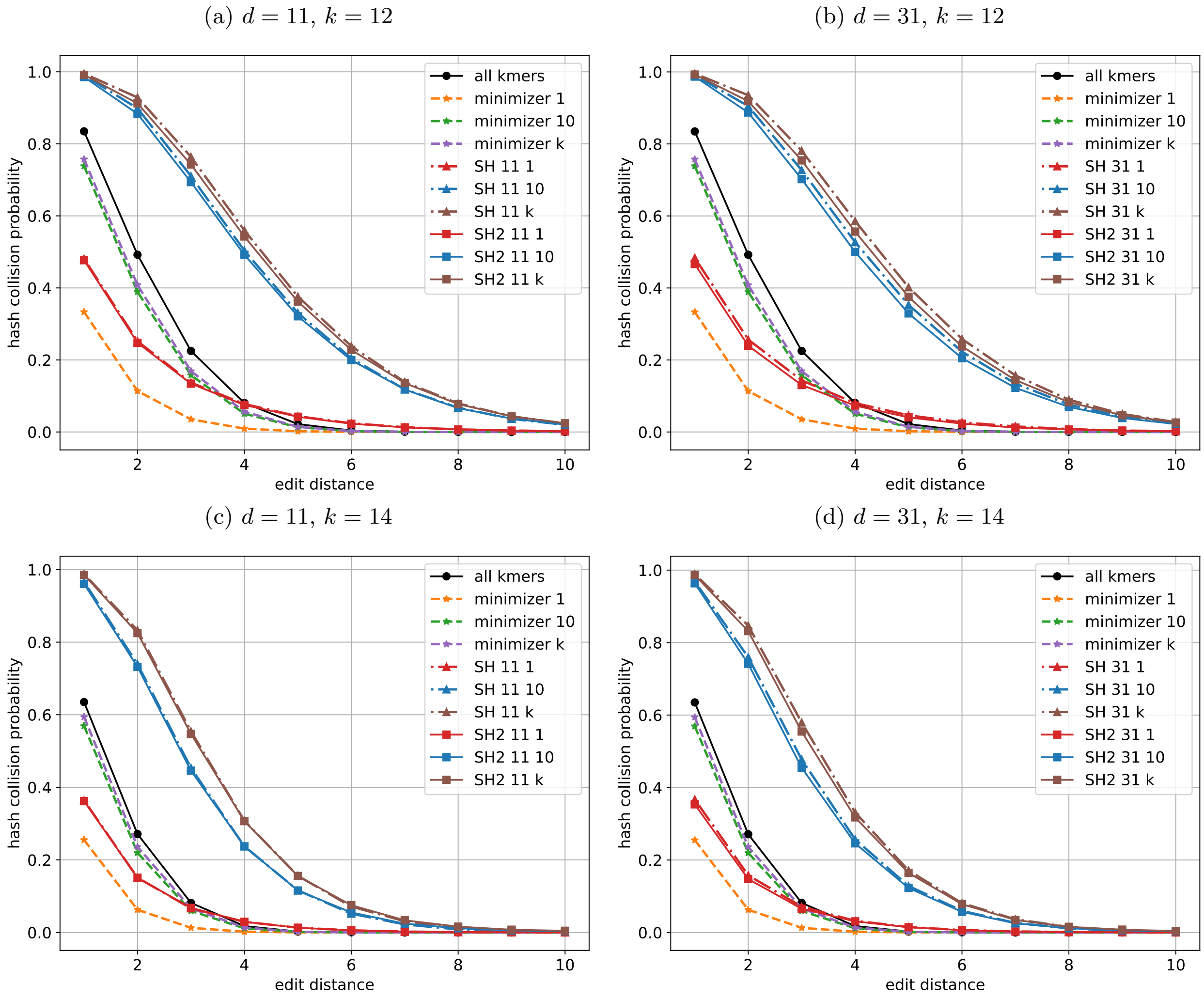

Figure 1: The probability of hash collision, estimated using simulations, for different seeding methods with $n = 20$ and $k = 12, 14$.

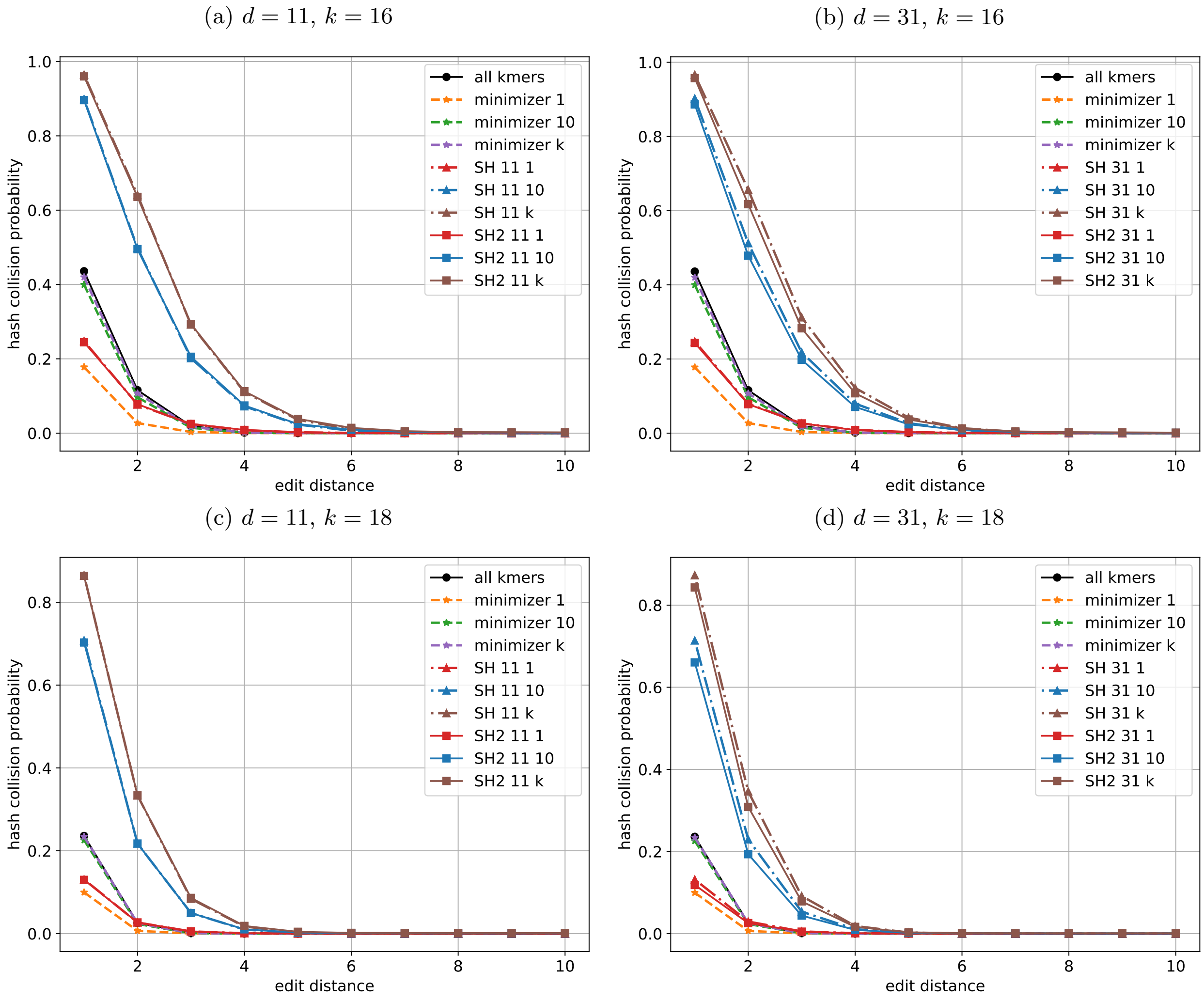

Figure 2: The probability of hash collision, estimated using simulations, for different seeding methods with $n = 20$ and $k = 16, 18$.

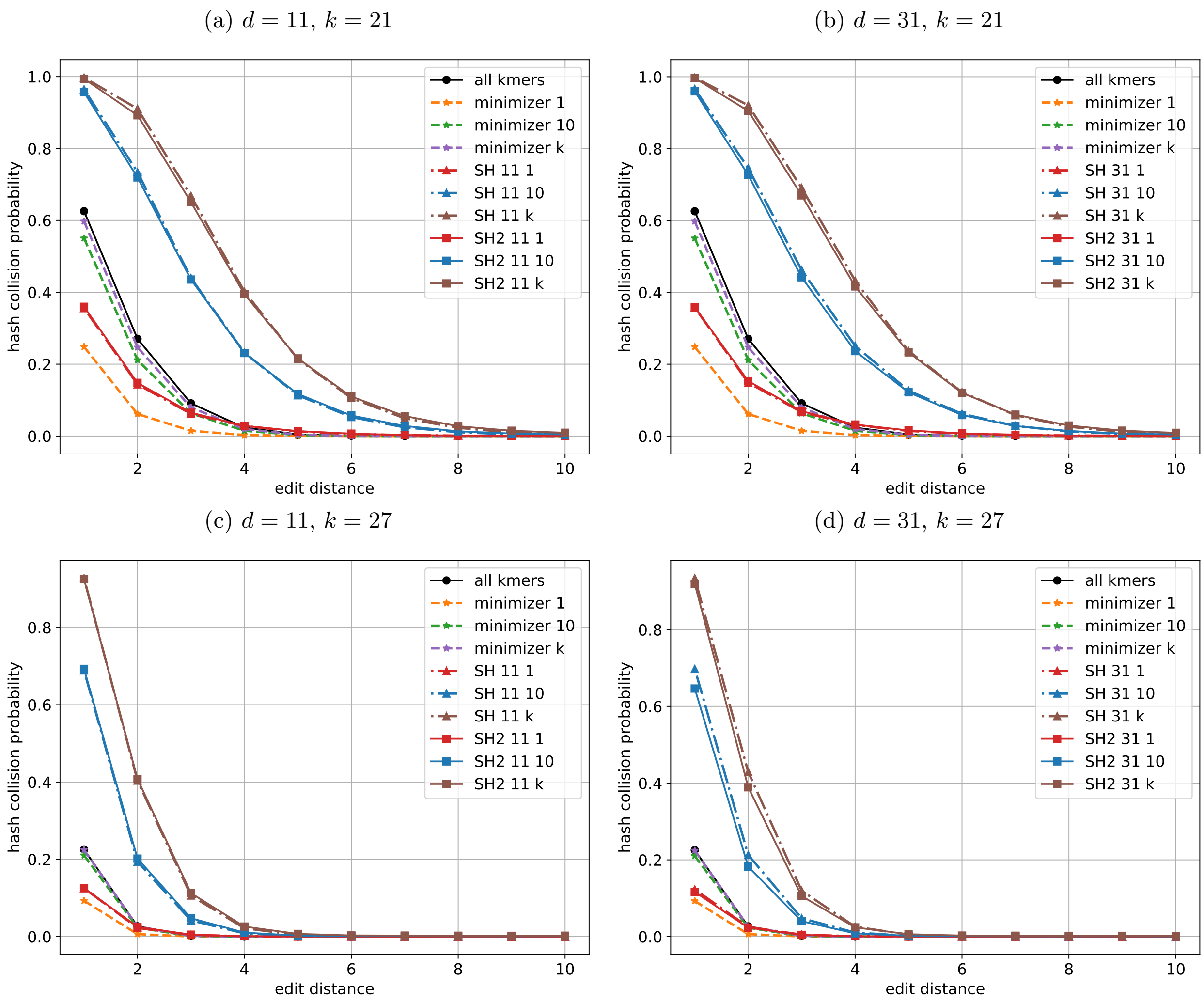

Figure 3: The probability of hash collision, estimated using simulations, for different seeding methods with $n = 30$ and $k = 21, 27$.

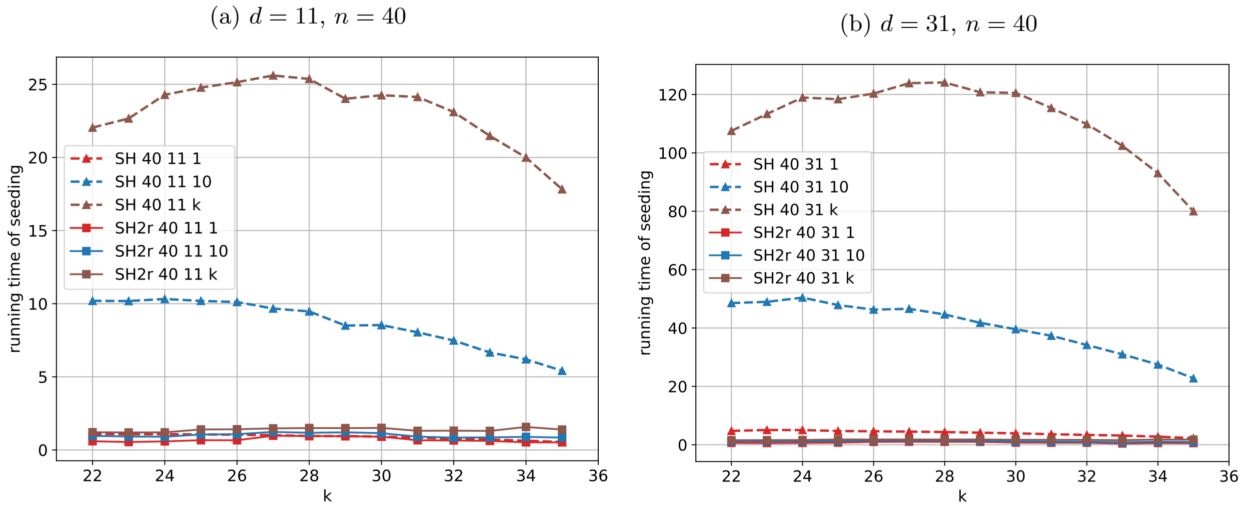

Figure 4: The average CPU time (second) of SubseqHash and SubseqHash2 per read with $n = 40$.

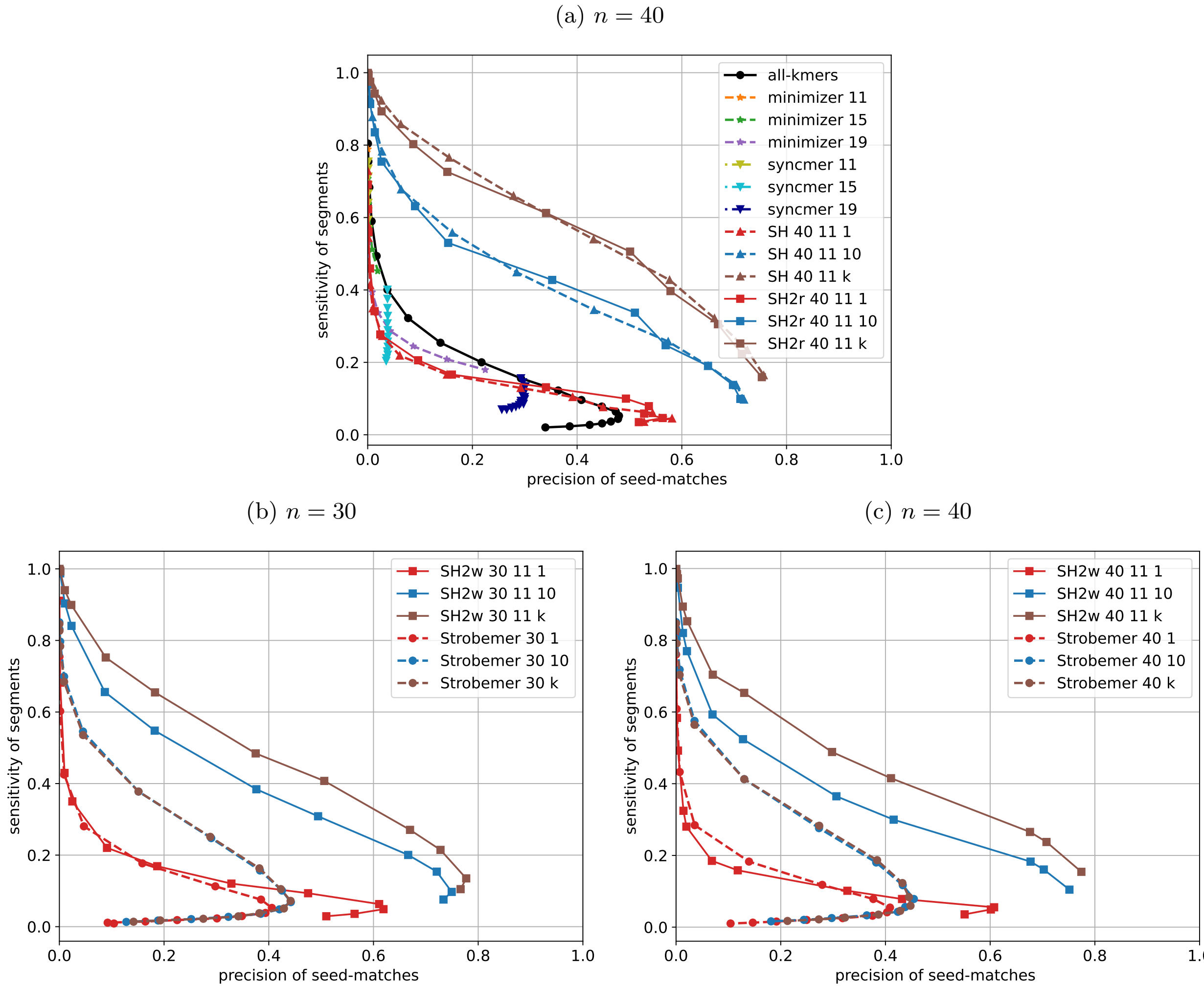

Figure 5: The average precision of seed-matches and sensitivity of segments with different seeding methods with $n = 40$.

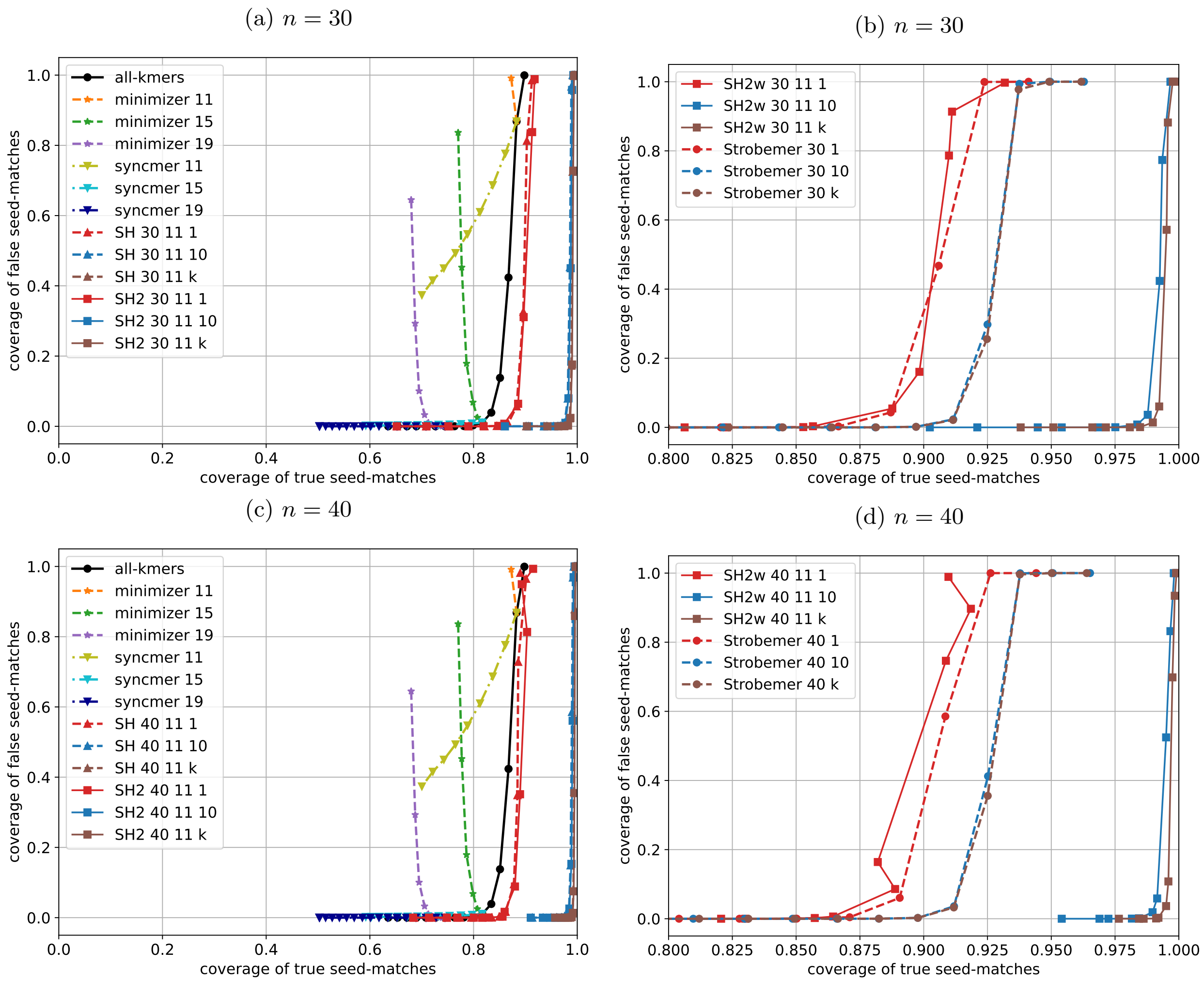

Figure 6: The true and false coverage of different seeding methods in sequence alignment with error rate $r = 5\%$, $n = 30, 40$. Figure (a) and (c) compare SubseqHash2 with all-kmers, minimizer, syncmer, and SubseqHash. Figure (b) and (d) compare SubseqHash2w with Strobemer.

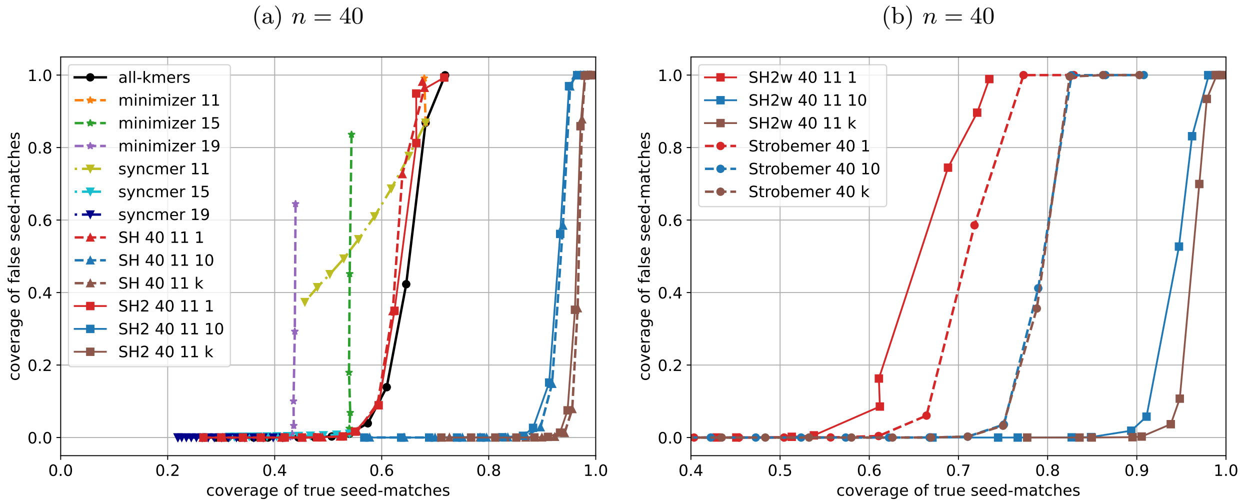

Figure 7: The true and false coverage of different seeding methods in sequence alignment with error rate $r$ = 10%, $n$ = 40. Figure (a) compares SubseqHash2 with all-kmers, minimizer, syncmer, and SubseqHash. Figure (b) compares SubseqHash2w with Strobemer.

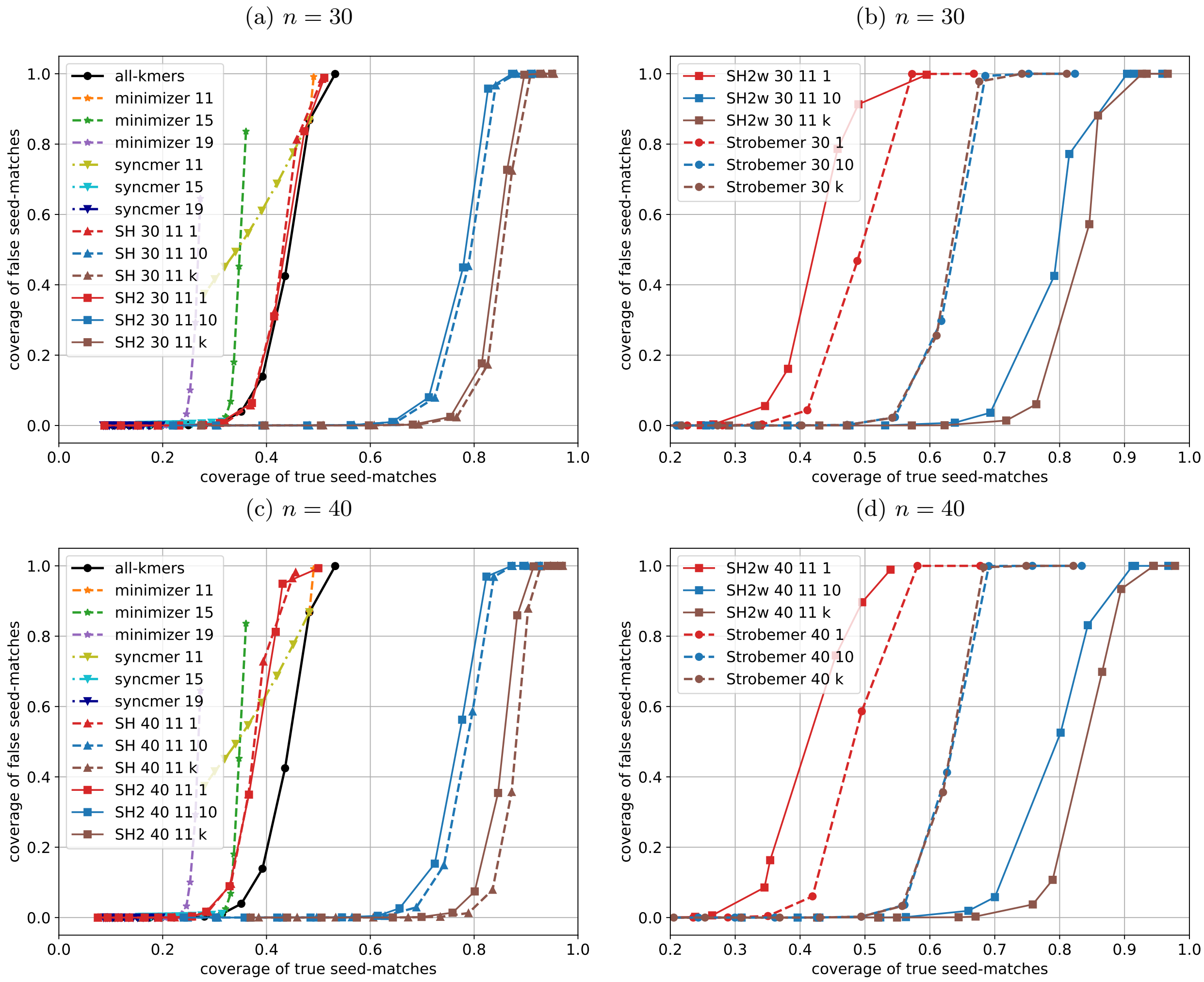

Figure 8: The true and false coverage of different seeding methods in sequence alignment with error rate $r = 15\%$, $n = 30$, $40$. Figure (a) and (c) compare SubseqHash2 with all-kmers, minimizer, syncmer, and SubseqHash. Figure (b) and (d) compare SubseqHash2w with Strobemer.

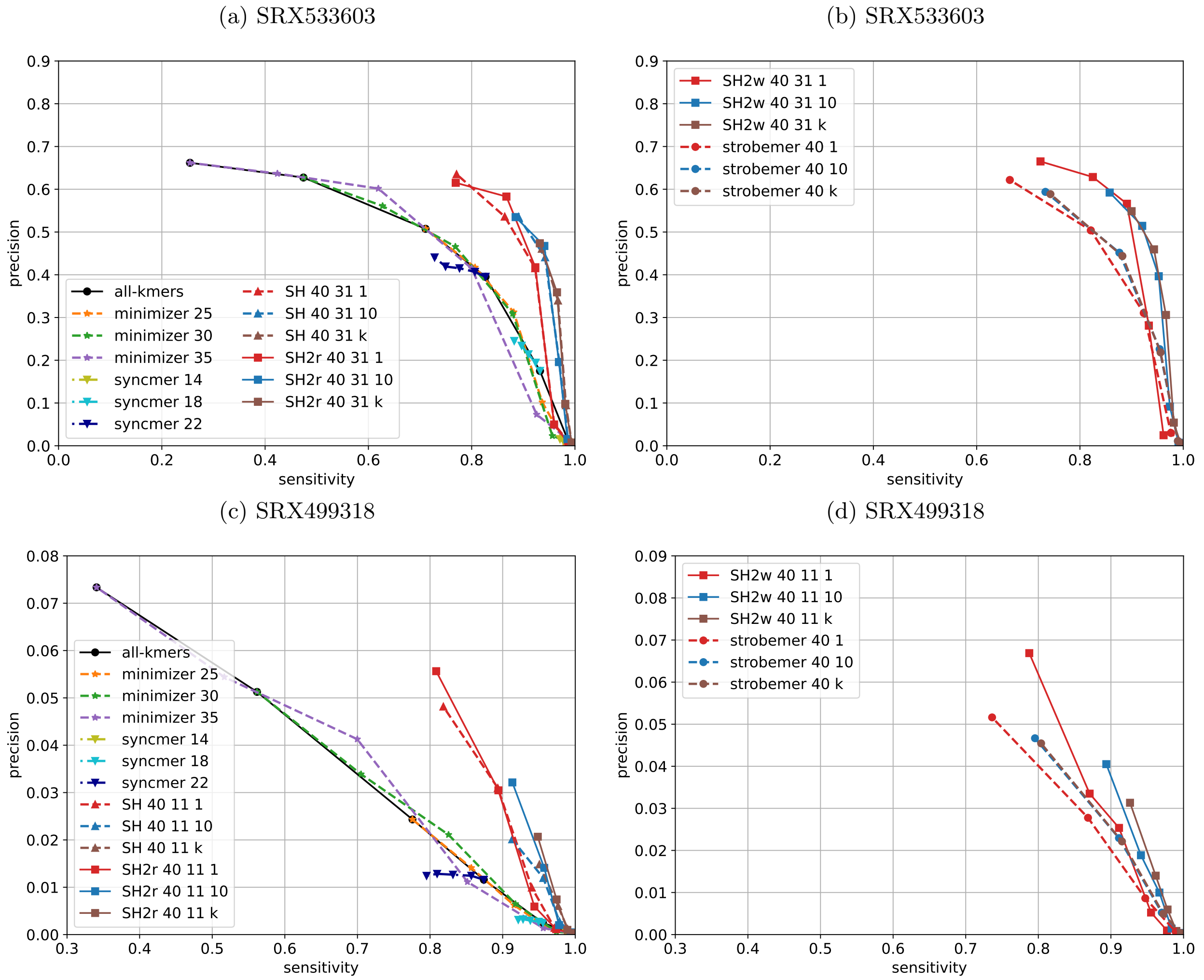

Figure 9: Overlap detection results on 10,000 reads sampled from the the *E. coli* SRX533603 and *D. melanogaster* SRX499318 dataset with $n = 40$. Figure (a) and (c) compare SubseqHash2r with all-kmers, minimizer, syncmer, and SubseqHash. Figure (b) and (d) compare SubseqHash2w with Strobemer.

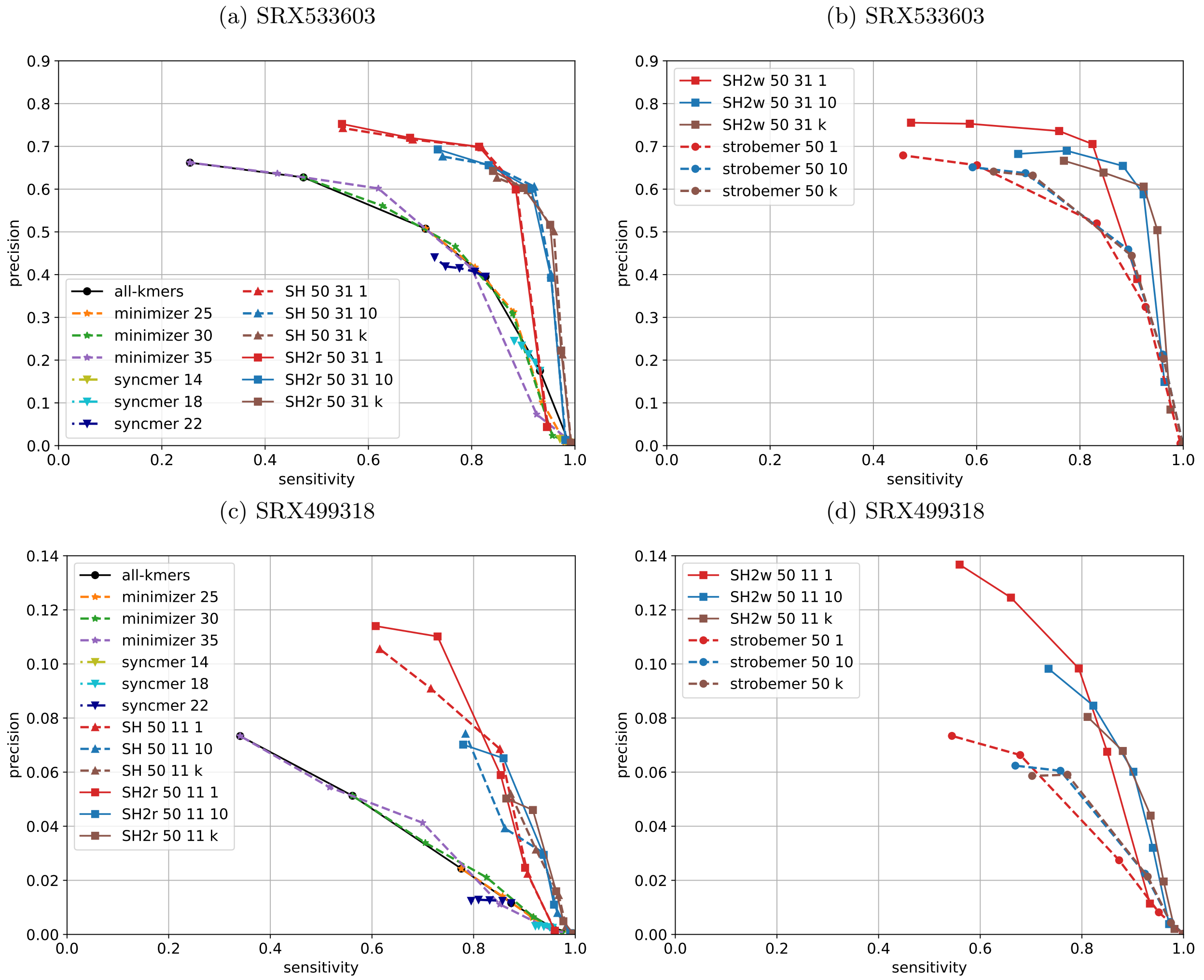

Figure 10: Overlap detection results on 10,000 reads sampled from the the *E. coli* SRX533603 and *D. melanogaster* SRX499318 dataset with $n = 50$. Figure (a) and (c) compare SubseqHash2r with all-kmers, minimizer, syncmer, and SubseqHash. Figure (b) and (d) compare SubseqHash2w with Strobemer.

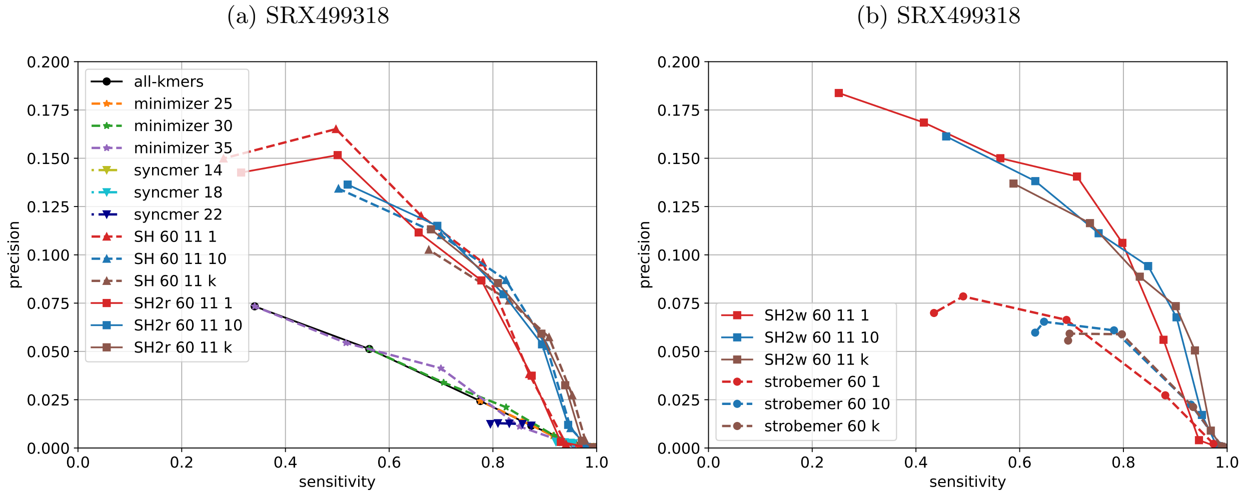

Figure 11: Overlap detection results on 10,000 reads sampled from the the *D. melanogaster* SRX499318 dataset with $n = 60$. Figure (a) compares SubseqHash2r with all-kmers, minimizer, syncmer, and SubseqHash. Figure (b) compares SubseqHash2w with Strobemer.
